# Network modularity reveals context and state-dependent reorganization of time-varying functional connectivity in single-cell resolved neural activity recordings

**DOI:** 10.1101/2025.11.02.686048

**Authors:** Sihoon Moon, Jingting Liang, Hyun Jee Lee, Ava Maalouf, Zikai Yu, Sahil Moza, Yun Zhang, Hang Lu

## Abstract

An important goal of neuroscience is to understand how biological neural networks organize activity at multiple scales to enable complex information processing and behavioral output. To address this challenge, large-scale neural activity datasets with increased resolution and wider coverage have become more prevalent across many model systems. However, bridging the gap in scale between changes in pairwise functional connectivity between neurons and changes in brain-wide organization of activity remains a key challenge. In this work, we demonstrate application of modularity-based community detection and network modularity to single-cell resolved recordings, for the first time, as a method to summarize complex changes in time-varying functional connectivity, facilitating comparisons across multiple time windows, recordings, and conditions. We apply these methods to both single-cell resolved multi-cell and whole-brain activity recordings. In the multi-cell recordings, we find that food odor changes functional connectivity between existing network modules in a *C. elegans* locomotory interneuron network, rather than reorganizing them. In spontaneous whole-brain activity, we identify several key hub neurons and combinations that significantly destabilize module assignments when silenced. Together, these results demonstrate community detection and modularity as a method for detecting context and network state-dependent changes in functional connectivity at the intermediate scale of network modules in single-cell resolved neural activity. Results from these analyses facilitate future investigation of mechanisms that mediate organization of neural activity at intermediate scales.

## Introduction

Advances in microscopy, image processing methods, and technology more broadly have increased the prevalence, scale, and resolution of neural activity data. Some notable examples include neuropixels arrays,^1^ the MICrONS dataset,^2^ and single-cell resolved neural activity.^3–8^ These datasets represent exciting advances as the combination of cellular resolution and wide-scale coverage promises the potential to link molecular, synaptic, and cellular level mechanisms with systems-level understanding of information processing by biological neural networks. This shared motivation of linking systems-level understanding to molecular and cellular pathways has also motivated recording single-cell resolved neural activity with brain-wide coverage in *Caenorhabditis elegans*. Comparatively, this field is relatively mature, with the first such demonstration in 2013^4^ and demonstration of freely-moving whole-brain imaging in 2015.^6,7^ However, recent advances in microscopy and image processing methods^9–14^ have made availability of such datasets increasingly prevalent.^15–22^

Despite experimental advances, analysis of time-varying functional connectivity in these datasets remains uncommon. Even for the compact neural network of *C. elegans* (302 neurons total, ∼100 – 180 in the head ganglion), adding the dimension of time makes direct inspection and interpretation of changes in pairwise functional connectivity intractable. In particular, some method of summarizing changes in functional connectivity across many time-windows, individual recordings, and experimental conditions is needed. Without these summaries, comparisons of how network structure changes across relevant experimental conditions (*e.g.* external stimuli, genetic or functional perturbations) are infeasible. Making such comparisons is essential for investigation of mechanisms that facilitate multi-scale organization of brain activity.

One possible approach to address this need is summarizing complex changes in time-varying functional connectivity using network modularity.^23^ Network modularity is a widely used method for detecting and evaluating community structure, which has been applied in many domains,^24,25^ including analysis of social networks,^26^ microbial ecology,^27^ metabolic networks,^28^ supply chains,^29^ shipping networks,^30^ and in neuroscience towards analysis of both anatomical and functional connectivity.^31–36^ Drawing inspiration from these examples, we apply modularity-based community detection to single-cell resolved neural activity recordings for the first time to summarize context and state-dependent changes in time-varying functional connectivity. Network modularity summarizes these changes by providing a metric that can be used to detect network modules (groups of neurons that connect strongly with each other), in addition to scoring how strongly connections are concentrated within detected modules.

Here, we apply these methods to two datasets: single-cell resolved neural activity data from a locomotory interneuron network (*glr-1* expressing neurons) during food odor stimulus and spontaneous whole-brain activity when key hub neurons are silenced.^17^ The compact summary of time-varying functional connectivity these methods provide enables comparison across multiple time windows, recordings, and conditions. By applying these methods, we find that food odor changes functional connectivity across existing modules but does not reorganize them. In contrast, network modules destabilize significantly when several key hub interneurons are silenced. These results connect changes in time-varying functional connectivity across different contexts (buffer versus food odor stimuli) and network states (silencing of different interneurons) to changes in network structure at the intermediate scale of network modules.

Bridging the gap between pairwise and brain-wide changes is increasingly relevant for large-scale recordings with both cellular resolution and brain-wide coverage, where a key challenge is uncovering mechanisms that enable multi-scale coordination of neural activity.

## Results

### Network modularity summarizes complex changes in time-varying functional connectivity

For single-cell resolved neural activity, especially in *C. elegans*, functional connectivity has been often analyzed under the assumption of time-invariance. While this assumption is convenient, collapsing the dimension of time in exchange for expanding the dimension of neurons into neuron-pairs, the limitations can be illustrated using a toy example, where functional connectivity between neurons changes between two different states (Figure 1a – c). While assuming time-invariance and calculating correlations across the full recording (Figure 1a, ii) correctly identifies the presence of strong connectivity between these neurons, it fails to identify the temporally localized nature of these strong connections to state 2 (Figure 1a, i 70 – 130 s). Calculating correlations within shorter windows can capture these state-dependent relationships between neurons (Figure 1b).

**Figure 1.**
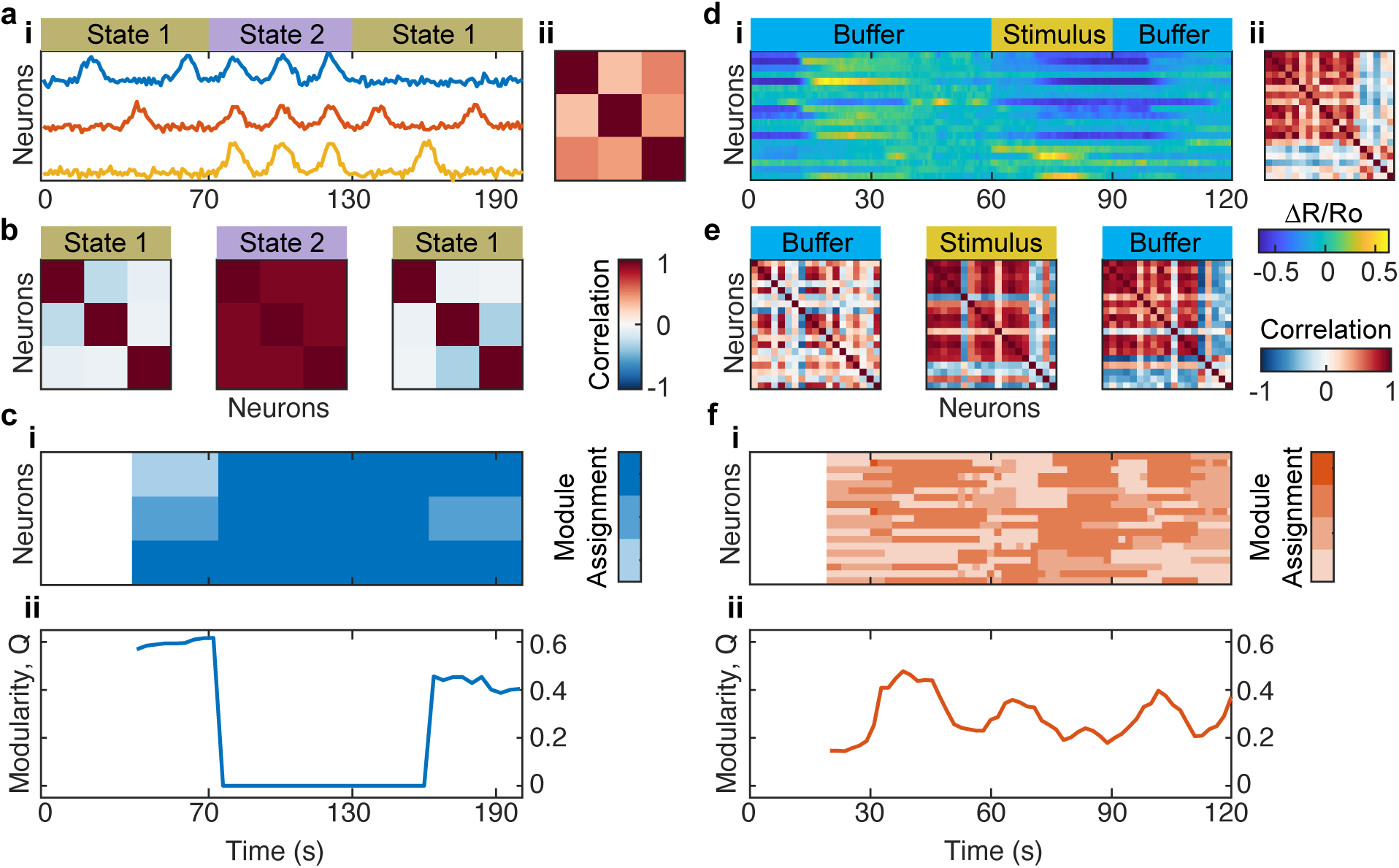
Network modularity summarizes complex changes in time-varying functional connectivity. **(a)** Neural activity (i) and time-invariant functional connectivity (ii, correlations across the full recording) for a toy example where functional connectivity is state dependent. **(b)** Calculating correlations for specific time windows (states) reveals temporally localized, state-dependent nature of strong correlations within the example circuit **(c)** Module assignments (i) and modularity (ii) summarize changes in time-varying functional connectivity (sliding window correlations), when specific timings of state transitions are unknown. **(d)** Multi-cell neural activity recording (ii) demonstrating both spontaneous (∼0-10s) and stimulus-driven (60-90s) neural activity and time-invariant functional connectivity. **(e)** Time-varying functional connectivity shows changes in network organization across different stimulus states. **(f)** Module assignments (i) and modularity (ii) show both spontaneous and stimulus-correlated (60-90s) fluctuations in modularity in real data.

In this toy example, directly inspecting and comparing correlation matrices is feasible because we know exactly when state transitions occur. This information lets us avoid sliding window correlations and only select a few appropriate windows to inspect. Unfortunately, we rarely have this information for real data. Even when external stimuli are applied, spontaneous internal dynamics may shift states and alter stimulus processing.^37^ As a result, it is favorable to use sliding windows to obtain time-varying functional connectivity for real data. However, even for this toy example with only three neurons, direct inspection of 10s or 100s of sliding window correlation matrices is infeasible, let alone for real recordings with 10s or 100s of neurons and an objective of comparing multiple windows, recordings, and conditions.

To address this, we demonstrate the application of community detection using network modularity to summarize complex changes in time-varying functional connectivity. These methods provide both a series of time-varying module assignments (Figure 1c, i) and corresponding modularity values (Figure 1c, ii). Applying these methods to our toy example, we see a decrease in network modularity when the network transitions to state 2 (Figure 1c, ii), suggesting potential reorganization. Inspection of the corresponding module assignments (Figure 1c, i) reveals the coalescence of the three previously independent neurons into a single module.

To demonstrate the utility of this analysis to a real dataset, we next apply it to an example *C. elegans* neural activity recording, where chemosensory stimulus has been applied using a microfluidic device (Figure 1d – f). These single-cell resolved multi-cell and whole-brain imaging datasets in *C. elegans* have become increasingly prevalent and are often collected under contexts where functional connectivity can change meaningfully over time (e.g. chemosensory stimulation with microfluidic devices,^16,38–40^ complex behaviors,^21,41^ and behavioral state transitions^15,42^). However, analysis of time-varying functional connectivity has remained uncommon. If analyses of time-varying functional connectivity in *C. elegans* became more common, they could be integrated with extensive knowledge of *C. elegans* neuroanatomy, gene expression, and functions of individual neurons to reveal mechanisms that organize functional connectivity at the network scale. When applied to our example recording (Figure 1d, i), which demonstrates both spontaneous (∼0 – 10 s) and stimulus-driven activity (60 – 90 s), we see changes in functional connectivity across different stimulus contexts (Figure 1e). Additionally, these context-specific connectivity matrices differ from the time-invariant functional connectivity (Figure 1d, ii). To summarize changes in functional connectivity over time across multiple windows, we apply community detection to obtain module assignments and calculate modularity values (Figure 1f). In this example, we see potential reorganization of modules across time (Figure 1f, i) and fluctuations in modularity (Figure 1f, ii).

### Food odor stimulus decreases network modularity in a small interneuron network

To demonstrate the efficacy of modularity-based community detection and network modularity for summarizing complex changes in time-varying functional connectivity, we apply these methods to a multi-cell food odor stimulus dataset. In this dataset, activity is recorded from a subset of neurons (*glr-1p*) that express GCaMP6s and nuclear localized mCherry, while applying food odor stimulus (*E. coli* OP50 culture supernatant) using a microfluidic device (Figure 2a). While limiting expression of fluorophores to a subset of neurons sacrifices brain-wide coverage, it improves accuracy and scalability of automated neuron tracking, trace extraction, and manual verification of the results. As a result, multi-cell imaging represents a more accessible and scalable use case, with greater applicability to biological problems where comparison across many genotypes, experimental conditions, and large sample sizes are needed. The specific subset of neurons recorded expresses a glutamate receptor subunit (*glr-1*) and includes several key command interneurons (e.g. RIM, AVA),^43^ which have been shown to encode reverse/forward locomotion states during spontaneous brain-wide activity. ^44^

**Figure 2.**
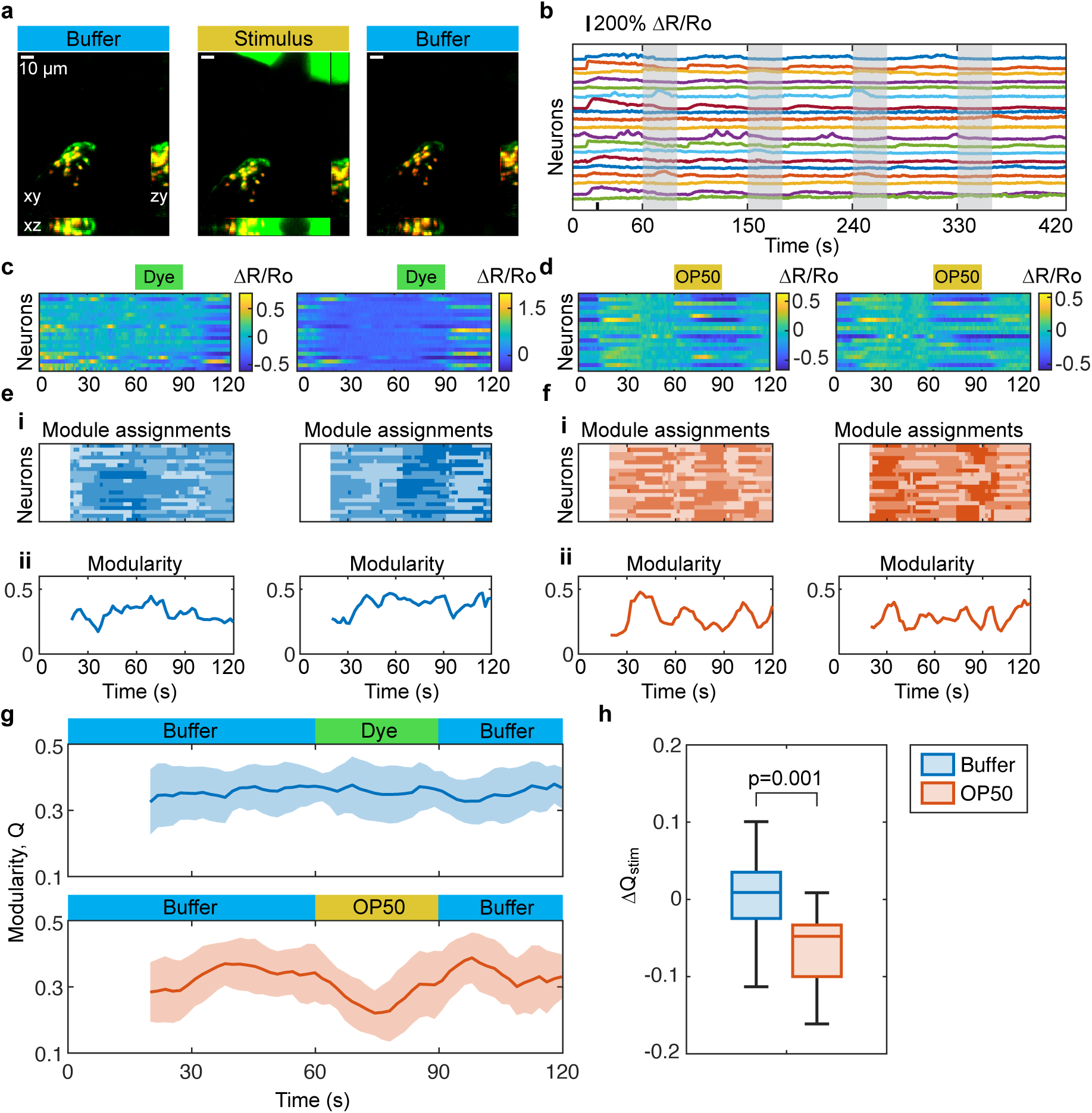
Food odor stimulus decreases network modularity in a small interneuron network. **(a)** Sample max projection images (xy, zy, and xz) from multi-cell imaging recording of *C. elegans* expressing GCaMP6s (green) and nuclear localized mCherry (red) in *glr-1* expressing neurons. During recording, stimulus (green) is delivered using a microfluidic device. Scale bars are 10 µm. **(b)** Neural activity vs. time for sample recording. Each *C. elegans* was stimulated 4 times for 30s each (gray bars) with 60 intervals in between. **(c, d)** Heatmaps of interneuron activity for individual stimulation trials show coordinated spontaneous activity during switches between odorless buffer (c) and stimulus correlated activity in response to application of *E. coli* OP50 food odor (d). **(e, f)** Module assignments (i) and modularity (ii) for individual trials shown in (c,d) demonstrate spontaneous fluctuations in modularity over time in both conditions (e and f, ii). **(g)** Average modularity decreases during food odor stimulus (bottom, 60-90s), but remains stable during buffer control condition (top, 60-90s). Shaded region corresponds to standard deviation across trials (n = 5 worms per condition x 4 trials per worm). **(h)** Decrease in modularity during OP50 stimulus application (ΔQ_stim_) is significant relative to ΔQ_stim_ during buffer control (p=0.001). Significance value obtained by fitting a linear mixed effect (LME) model that controlled for repeated measurements and potential trends across trials (methods).

In the context of chemotaxis behavior, the network should switch to the forward state when food odor is applied and reversals when removed. These switches allow the animal to continue towards a food lawn when leaving and return to the lawn when exiting. As a result, we see coordinated activation and inhibition of several neurons when applying a food odor stimulus (*E. coli* OP50 supernatant) using a microfluidic device (Figure 2a, b, d). We show two example trials of stimulus application that demonstrate this coordinated activation and inhibition in figure 2d. For comparison, we also show spontaneous activity for two trials, where the “stimulus” is a switch between odorless buffer and odorless fluorescent dye instead of food odor (Figure 2c). In addition to measuring spontaneous activity, the buffer condition controls for inadvertent mechanical and thermal stimulation by the microfluidic device, sensation of dye rather than the food odor stimulus, and how these might affect module assignments and modularity. For both the spontaneous (Figure 2e) and OP50 stimulus (Figure 2f) example recordings, we see changes in module assignments and modularity over time.

While modularity fluctuates over time for individual trials of spontaneous activity (Figure 2e, ii), these fluctuations are not synchronized to specific time periods. As a result, average modularity across trials is constant across time (Figure 2g, top). In contrast, average modularity across trials decreases when applying OP50 food odor (Figure 2g, bottom, 60 – 90s). We quantify these decreases by averaging the modularity during stimulus (60 – 90 s) and non-stimulus (20 – 60 s, 90 – 120 s) periods and taking the difference, Δ*Q_stim_* (Figure 2h). Comparing Δ*Q_stim_* between OP50 and buffer conditions, we see a significant decrease for OP50 (p = 0.001). This significance value was obtained by fitting a linear mixed effect (LME) model that accounts for repeated observation of individuals and potential trends across trials.

These consistent decreases in modularity during food odor stimulus across multiple worms and trials suggest some form of network reorganization, either reorganization of network modules or changes in functional connectivity between existing modules. While further analysis is needed to suggest the exact nature of the changes occurring (Figure 3), modularity has provided a compact metric for summarizing, aggregating, and detecting these changes across multiple time windows, trials, individuals, and conditions. This summary is especially valuable for this circuit, where spontaneous activity and variable timing of activity relative to stimulus^37^ requires use of sliding windows, rather than selection of specific windows.

**Figure 3.**
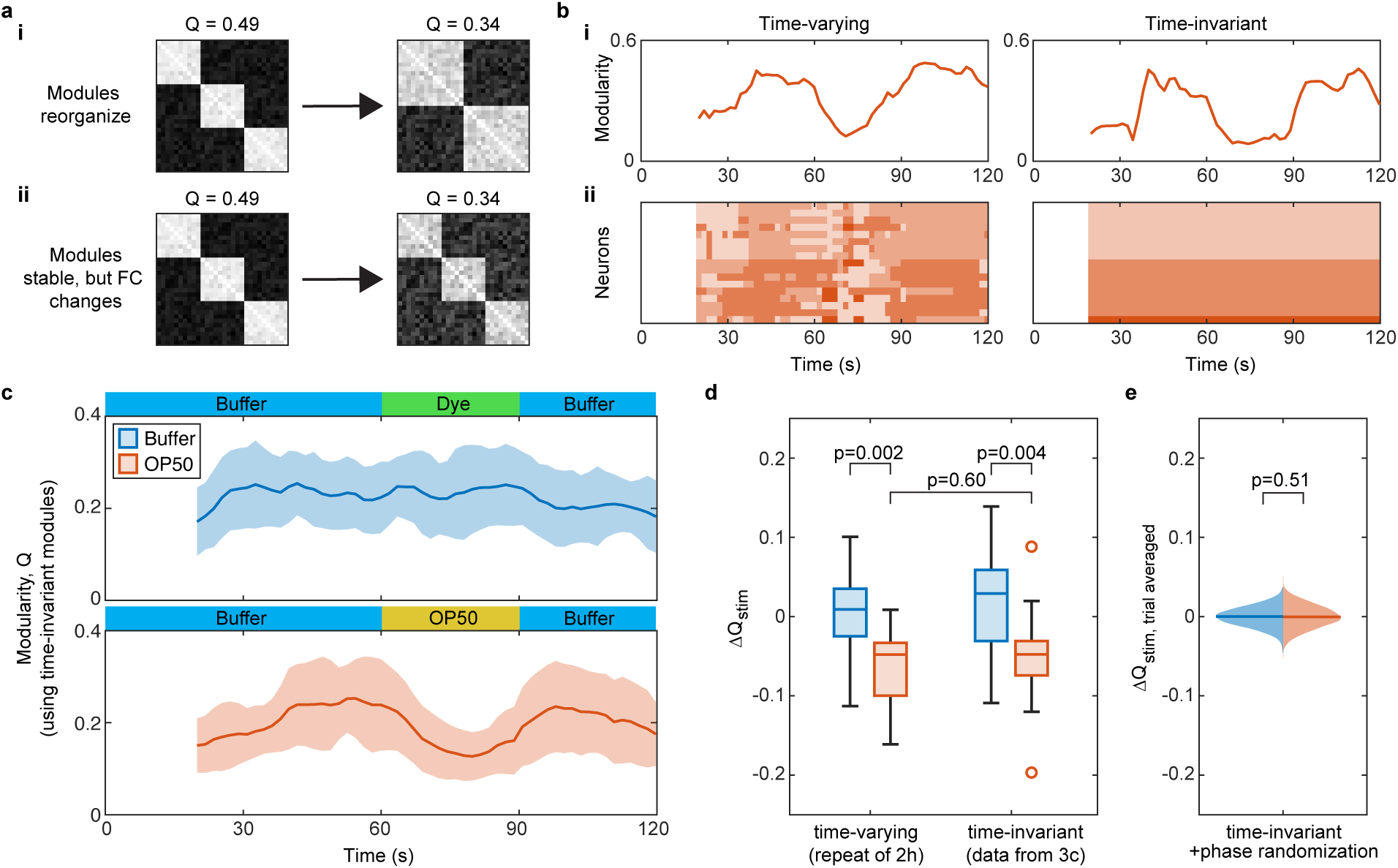
Changes in correlation strength between modules, instead of module reorganization, drives decreased modularity during food odor stimulus. **(a)** Similar changes in network modularity can results from different changes in underlying network structure. A decrease in modularity could either result from reorganization of network modules (i) or stable module organization, but changes in functional connectivity (FC) between modules. **(b)** To test effect of network reorganization on changes in modularity, we recalculate modularity (i) using time-invariant module assignments (ii, right) and compare the results obtained using time-varying (left) and time-invariant module assignments (right). Sample trial shown is from the food odor stimulus condition. **(c)** Modularity still changes in response to food odor (60-90s, bottom) when calculated with time-invariant modules. Lines are average modularity and standard deviation across trials (shaded region) for buffer control (top) and food odor stimulus (bottom) conditions, calculated using time-invariant modules. Axes limits differ from Figure 2g to facilitate visualization. **(d)** Decrease in modularity during food odor stimulus application ΔQ_stim_ is still significant when calculated using time-invariant modules (p=0.004) and value of food odor response does not differ significantly when calculated with time-varying module assignments (p=0.60). Significance values obtained by fitting an LME model that accounts for repeated measurements of individual worms (n = 5 worms per condition x 4 trials per worm). **(e)** Recalculation of modularity and with time-invariant modules and phase randomized correlations (n=1,000) eliminates condition-averaged decrease in modularity during food odor stimulus application (p=0.51). Significance corresponds to fraction of phase randomizations where condition averaged food odor stimulus was greater for OP50 than buffer (methods).

### Changes in correlation strength between modules, instead of module reorganization, drives decreased modularity during food odor stimulus

While changes in network modularity reveal some form of change in network structure is occurring, they do not reveal their exact nature (Figure 3a). Similar changes in could result from either reorganization of network modules (change in size/number of modules, Figure 3a, i), or changes in functional connectivity between existing modules, without reorganization (Figure 3a, ii). To evaluate how strongly reorganization might influence decreases in modularity during food odor application, we recalculated modularity using time-invariant module assignments (Figure 3b). Time-invariant module assignments were obtained by taking the consensus module assignment across all timepoints, then applying this consensus assignment to each timepoint (Figure 3b, ii) to recalculate modularity (Figure 3b, i). Even when module assignments are time-invariant, changes in functional connectivity across time will lead to changes in modularity.

When modularity is recalculated using these time-invariant module assignments, we still observe consistent stimulus-correlated decreases in modularity (Figure 3c, bottom, 60-90s). We tested the effect of calculating modularity using time-invariant modules by fitting a linear mixed effects (LME) model. The decrease in modularity during 60-90s (Δ*Q_stim_*) is still significant for the OP50 food odor condition, relative to the decrease observed for buffer control (p = 0.004, Figure 3d), when calculated with time-invariant modules. Additionally, there was no difference in Δ*Q_stim_* for the OP50 condition when calculated with time-invariant versus time-varying modules (p = 0.60). These results suggest that food odor changes strength of functional connectivity between network modules, rather than reorganizing network modules within the *glr-1p* command interneuron network.

One possible interpretation of this result, that time-invariant modules do not decrease sensitivity of modularity to food odor relative to time-varying, is that some of the changes in time-varying module assignments might reflect degeneracy in module assignments, rather than genuine reorganization. The potential for degeneracy, where dissimilar module assignments can produce similar, near-optimal modularity values, has been noted in previous literature.^45^ In light of this, the approach demonstrated here, which compares results obtained using time-varying and time-invariant module assignments, is an important control to account for this possibility.

To validate the role of changes in functional connectivity in driving stimulus-correlated changes in modularity, we re-calculated network modularity using a time-invariant modules and joint phase randomized functional connectivity (Figure 3e). The joint phase randomized functional connectivity preserves autocorrelation and frequency structure of time-varying functional connectivity for individual pairs of neurons, but randomizes phases of different frequency bands, eliminating meaningful changes in functional connectivity over time. By using the same set of randomized phases across all neuron pairs (joint phase randomization), we also preserve covariances of functional connectivity between neuron pairs across time. For each randomization, we calculated average Δ*Q_stim_* for each condition (Buffer, OP50) across all worms and trials. Across 1,000 randomizations, median trial averaged Δ*Q_stim_* was close to 0 for both conditions (Figure 3e). Additionally, trial-averaged Δ*Q_stim_* was equally likely to be lower for Buffer versus OP50 rather than higher (p = 0.51 across 1,000 randomization). The elimination of OP50 response only by phase randomization of time-varying functional connectivity, and not by substitution of time-varying modules with time-invariant, confirms that food odor alters functional connectivity within this network by changing connectivity between existing modules in the network, rather than reorganizing modules.

### Optimization of method and parameters to increase sensitivity of modularity to food odor

The results shown in figures 2 and 3 were obtained using a version of modularity that accounts for correlation signs^46^ and optimizing module assignments for individual time windows (single-slice). These methods were selected after comparison against the original modularity equation^23^ which only considers correlation strength (absolute value) (Figure 4a – b) and multi-slice modularity (Figure 4c – e), a modification of modularity-based community detection which couples adjacent timepoints to encourage greater temporal consistency of module assignments^47^.

**Figure 4.**
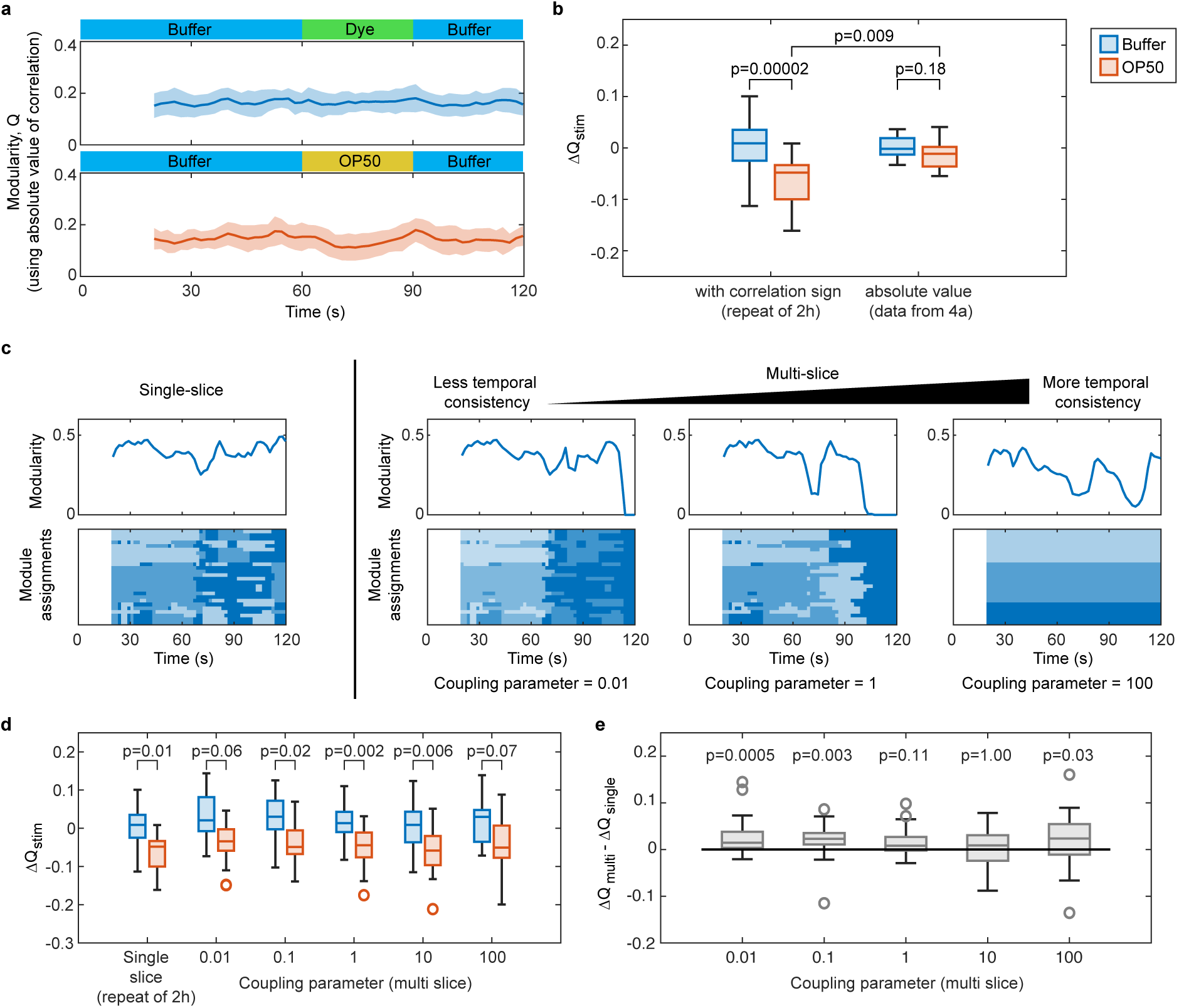
Optimization of method and parameters to increase sensitivity of modularity to food odor stimulus. **(a)** Modularity does not change significantly in response to food odor stimulus (60-90s, bottom) when calculated with absolute value of correlation. Lines and shaded regions correspond to average modularity and standard deviation across trials for buffer (top) and OP50 (bottom) conditions, after reoptimizing modules and calculating modularity with absolute value. Axes limits adjusted to match Figure 3c. **(b)** Decrease in modularity during stimulus application ΔQ_stim_ no longer significantly differs between OP50 buffer conditions (p=0.18) when modularity is calculated using absolute value and value of OP50 response differs significantly between methods (p=0.009). **(c)** Comparison of modularity and module assignments obtained for a sample spontaneous activity recording using single-slice (left) and multi-slice module assignment (right). For multi-slice module assignment, increasing coupling parameter increases temporal consistency until module assignments become time-invariant (coupling parameter ∼100 in this example). Rather than converge towards single-slice, multi-slice module assignment results become more degenerate, resulting in some timepoints where all neurons collapse into a single module. **(d)** Difference in ΔQ_stim_ between buffer (blue) and OP50 (red) conditions is relatively insensitive to method used or value of coupling parameter selected, with only extreme values of coupling parameter (0.01 or 100) showing non-significant differences after Bonferroni correction. **(e)** Comparison of ΔQ_stim_ calculated using single-slice and multi-slice methods shows results are the most similar for this dataset when coupling parameter is 10 (p = 1.00, after Bonferroni correction and truncation), which balances degeneracy at low coupling parameter values with temporal-invariance imposed by strong coupling. Significance in b, d, and e were calculated by fitting LME models that accounted for repeated observations (n = 5 worms per condition x 4 trials per worm), calculation of pairwise contrasts, and Bonferroni corrected for multiple comparisons in d and e (n=6 comparisons in d, n=5 comparisons in e).

Inclusion of correlations signs is particularly relevant for single-cell resolved functional connectivity in *C. elegans,* which has been noted to demonstrate both strong negative and positive correlations, a feature that differs from the ranges of correlation values typically observed in fMRI datasets.^17,31,48^ When we ignore the excitatory/inhibitory nature of connections between neurons, reoptimize modules assignments, and calculate modularity based on absolute value of correlation alone, we no longer observe a decrease in modularity during OP50 stimulus (Figure 4a, 60 – 90 s). When correlation signs are ignored, the decrease in modularity during 60 – 90 s (Δ*Q_stim_*) is no longer significant for the OP50 food odor condition, relative to the change observed for the buffer control (p = 0.18, Figure 4b). Additionally, the observed Δ*Q_stim_* for OP50 differs significantly when calculated with absolute values instead of accounting for correlation signs (p = 0.009, Figure 4b). In addition to being useful for method optimization, this result also suggests that changes in excitatory/inhibitory relationships between neurons encode changes in network structure during food odor stimulus, rather than changes in correlation strength alone.

We also tested the effect of jointly optimizing module assignments across all time windows using multi-slice modularity^47^ instead of optimizing module assignments for individual windows (single-slice). Multi-slice modularity introduces a coupling parameter, ⍵, that encourages assignments of the same neuron to the same module across adjacent windows. The value of this coupling parameter defines how much the multi-slice modularity objective function benefits from keeping neurons in the same module across time. Increasing ⍵ increases temporal consistency, resulting in time-invariant module assignments when ⍵ is sufficiently high (Figure 4c). For testing multi-slice module assignment, we swept coupling parameter across a wide range of values (0.01, 0.1, 1, 10, 100) and followed previous guidelines for obtaining a representative consensus assignment across many optimization runs (n = 1,000).^34^ We expected that decreasing ⍵ (coupling parameter) would eventually result in comparable module assignments and modularity to single-slice (Figure 4c). However, converting local optimization of module assignments for individual windows to global optimization across all windows also increases degeneracy (instability of module assignments across optimization runs). This degeneracy results in some windows where all neurons collapse into a single module after obtaining a representative consensus assignment. This collapse is likely more prevalent for lower values of coupling parameter (0.01, 1 for the example recording shown in Figure 4c), due to the increased degeneracy that results from the increased number of communities detected across time with weaker coupling parameter values.

While module assignments diverge between multi-slice and single-slice methods, comparing the observed decrease in modularity during 60 – 90 s (Δ*Q_stim_*) for OP50 versus buffer across different methods (single-slice and multi-slice using different coupling parameter values) revealed that our results are relatively insensitive to selection of method or parameter values (Figure 4d). Only extreme values of coupling parameter (0.01, 100) resulted in non-significant differences between conditions after Bonferroni correction (Figure 4d). To determine which value of coupling parameter produced the results most similar to single-slice results, we took the difference in Δ*Q_stim_* observed using multi-slice and single-slice methods (Δ*Q_multi_* − Δ*Q_sin_*_(*le*_, Figure 4e). Lower coupling parameter values (0.01 – 1), where greater instability of module assignments across optimization runs is observed, generate skewed predictions (Figure 4e) relative to the single-slice results. Predictions from the multi-slice method are most representative of single-slice method results when coupling parameter is appropriately tuned (high enough to stabilize assignments across time, but low enough that assignments do not become time-invariant) (Figure 4e, coupling parameter = 10). These results re-emphasize the importance of appropriate coupling parameter tuning for multi-slice module assignment and suggest that single-slice module assignment (with re-indexing of individual timepoints to improve temporal consistency) can provide a valuable, non-user biased estimate of what results multi-slice might provide after appropriate parameter-tuning.

### Analysis of time-varying whole-brain functional connectivity reveals destabilization of modularity when key hub neurons are silenced

In the multi-cell food odor response dataset, external stimuli influenced changes in time-varying functional connectivity. However, functional connectivity can also vary over time due to spontaneous activity. We demonstrate the efficacy of modularity in summarizing changes in network structure in this context by applying modularity-based community detection to a spontaneous whole-brain imaging dataset from Uzel, Kato, and Zimmer 2022.^17^ The dataset contains spontaneous whole-brain activity for transgenic strains where activity of different key hub interneurons were silenced by targeted expression of a histamine gated chloride channel,^49^ in addition to wild-type spontaneous whole-brain activity. The authors identified these key hub interneurons after analyzing topological features of the *C. elegans* connectome and observed significant decorrelation of time-invariant functional connectivity after silencing.

To ask how these perturbations might affect time-varying functional connectivity, we assign modules and calculate modularity based on both correlation strength and signs, using single-slice module assignment. We use single-slice, rather than multi-slice, to compare different silencing conditions after testing multi-slice and observing that silencing different neurons results in different sensitivity to multi-slice coupling parameter (Supplementary Figures 1 – 5). In wild-type data, where no neurons have been silenced, modularity is relatively stable across time, although occasional decreases occur (Figure 5a, ∼400 s). When these decreases occur, several neurons change module assignments, but module assignments and modularity return to baseline. As a result, the wild-type allegiance matrix shows high modular allegiance between neurons (frequent co-clustering between neurons in the same module), indicating stable recruitment patterns across time. When three key hub interneurons involved in both forward and reverse locomotion (AVB, RIB, and AIB) are silenced, modularity becomes less stable (Figure 5b). Instead of two reliably recruited modules, like we see in wild-type, module assignments are destabilized, leading to decreased modular allegiance (Figure 5b).

**Figure 5.**
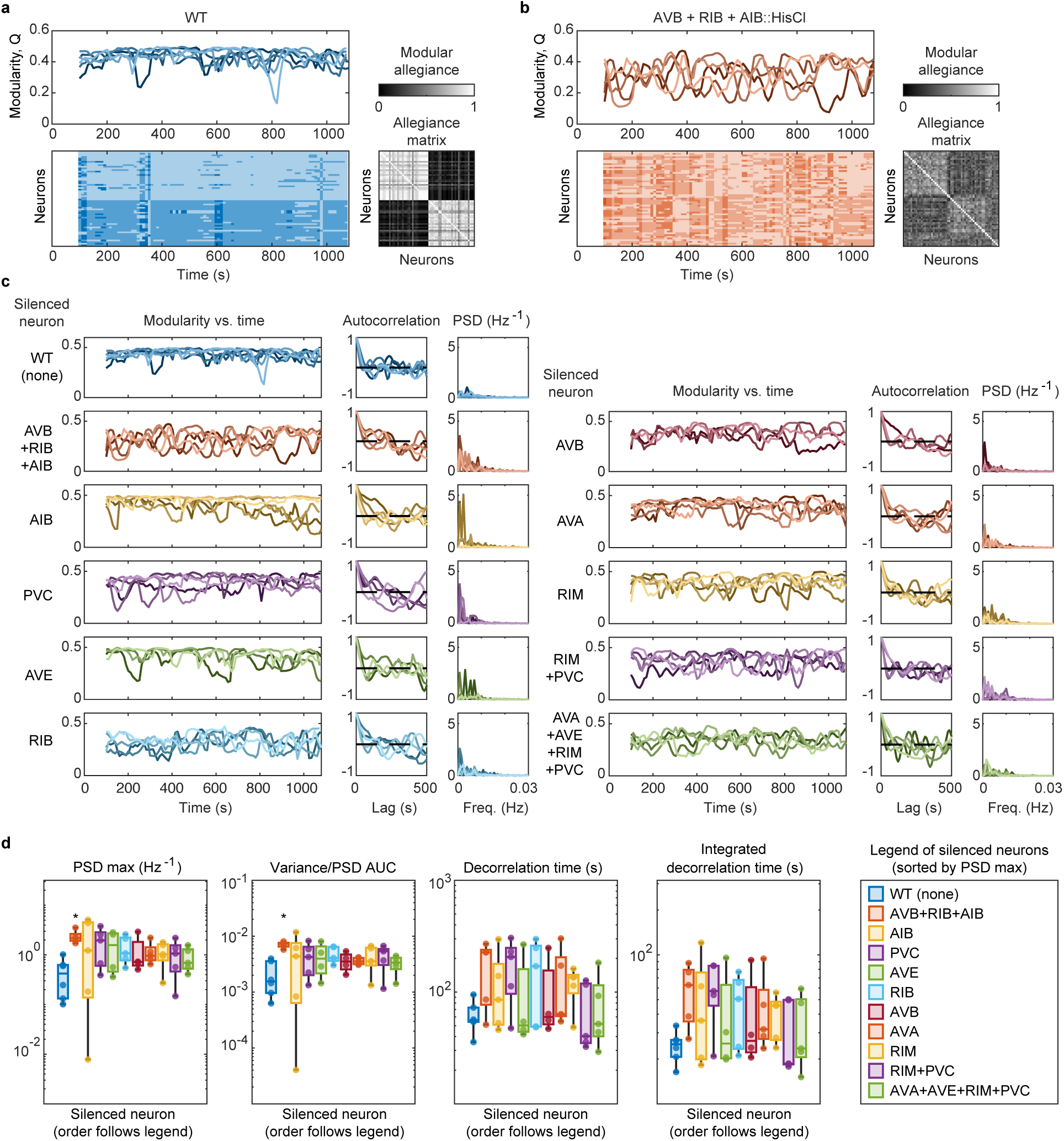
Analysis of time-varying whole-brain functional connectivity reveals destabilization of modularity when key hub neurons are silenced. **(a)** For wild-type (WT) spontaneous whole-brain activity recordings, modularity vs. time (top, n = 6) and module assignments for representative recording (bottom left) are relatively stable across time. Stability of module assignments across time can also be visualized using an allegiance matrix (probabilities of module co-assignment between neuron pairs), where two clear modules are present with reliable co-assignment (high modular allegiance) within modules and infrequent co-assignment (low modular allegiance) across modules. **(b)** When three key hub neurons are silenced (AVB + RIB + AIB::HisCl), modularity becomes less stable over time (top, n = 5) and module assignments become less stable (bottom left), which is also shown by more intermediate values of modular allegiance in the allegiance matrix (less reliable patterns of module co-assignment between neurons). **(c)** Modularity vs. time for WT spontaneous whole-brain activity is relatively stable, with autocorrelation quickly decaying to 0 and low power spectral density (PSD) values. Silencing activity of key hub interneurons destabilizes modularity vs. time, slowing decay of autocorrelations and increasing PSD (higher variance). After WT, silenced neurons are arranged from most destabilization to least (highest to lowest average PSD max) from top to bottom, column 1 to 2 (AVB+RIB+AIB silencing induces the most destabilization). **(d)** Quantifying destabilization as either max value of modularity PSD (left) or total variance (right) shows significant destabilization of modularity when AVB+RIB+AIB are silenced (p=0.03 left, p=0.04 right) but increase in integrated decorrelation time and decorrelation time are not significant (p=0.17 for integrated decorrelation time and p=0.51 for decorrelation time). Statistical significance was tested using pairwise Welch’s t-tests on log transformed stability metric values with Bonferroni correction for multiple comparisons.

We summarize the instability in modularity over time across all different neuron silencing conditions by plotting modularity, autocorrelation, and power spectral density (Figure 5c). The autocorrelation shows how quickly modularity samples its overall variance across time. If variance is primarily explained by high frequency noise, autocorrelation will quickly decay to zero, but if there is additional instability it will delay the decay of autocorrelation. Power spectral density (PSD) measures strength of oscillations in modularity relative to its average value, with greater values indicating greater instability. As can be seen in Figure 5c, wild-type modularity is relatively stable across time, with autocorrelation decaying relatively quickly and relatively low PSD values. By comparison, when AVB+RIB+AIB are silenced, modularity is less stable across time, autocorrelation decays more slowly, and higher PSD values are observed. The remaining neuron silencing conditions are sorted from most to least destabilization of modularity (max value of PSD across all frequencies, averaged across recordings per condition).

Compared to wild-type, silencing AVB+RIB+AIB significantly destabilizes modularity when we quantify stability as the max value of PSD across all frequencies (PSD max) or the variance of modularity over time (Figure 5d). Variance is also equal to area under the curve (AUC) of PSD across all frequencies. Decorrelation time (when autocorrelation first crosses zero) and integrated decorrelation time (AUC of autocorrelation before decorrelation time) also suggest increased instability when silencing these neurons, but the changes are not statistically significant. In contrast to silencing AVB+RIB+AIB in combination, the effect of silencing other neurons is more subtle. This significant destabilization of modularity when AVB+RIB+AIB are silenced might come from slight reduction of activity levels in this condition, with individual neurons demonstrating bouts of transient, unstable activation.^17^ Behaviorally, silencing these neurons in combination results in significant reduction in forward locomotion speed but no change in frequency of reversals.^17^

### Silencing key hub interneurons significantly destabilizes module assignments

While triple-silencing AVB, RIB, and AIB significantly destabilized modularity, destabilization when silencing other key hub neurons was minimal (Figure 5d). To test whether the same would hold true for stability of module assignments, we recalculated modularity for each recording using time-invariant module assignments (Figure 6b) and compared the results against those obtained using time-varying module assignments (Figure 6a). The stability of wild-type module assignments across time (Figure 6a, i) results in relatively little change in modularity when recalculated with time-invariant module assignments (Figure 6b, i). By contrast, recordings collected when AVB+RIB+AIB have been silenced are much more sensitive to recalculating modularity with time-invariant module assignments (Figure 6b, ii). This is reasonable given the instability of time-varying module assignments when these neurons are silenced (Figure 6a, i) and suggests that silencing these neurons destabilizes modularity by destabilizing module assignments across time.

**Figure 6.**
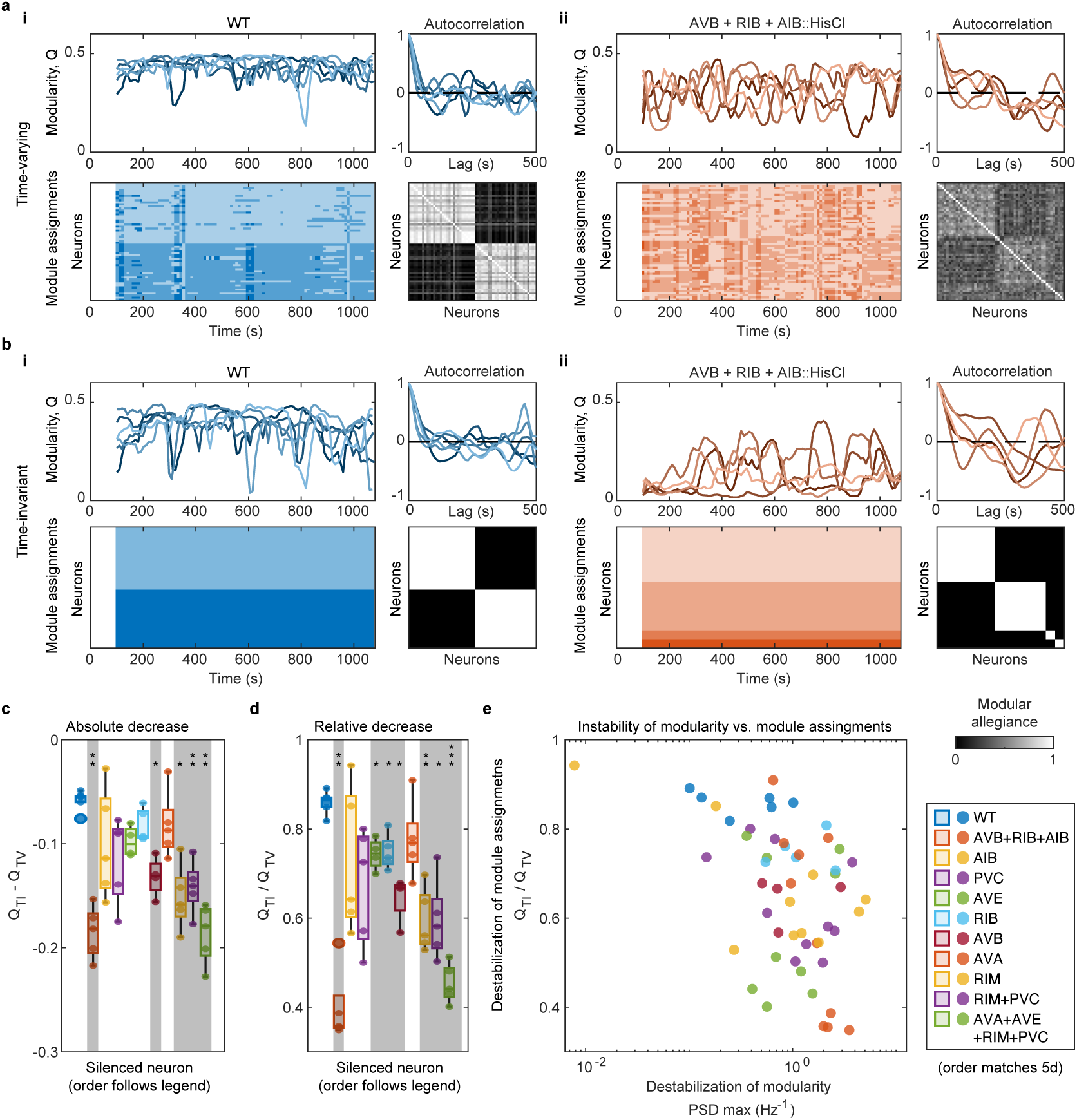
Silencing key hub interneurons significantly destabilizes module assignments. **(a, b)** Modularity vs. time, autocorrelation, module assignments, and allegiance matrix for WT (i) and AVB+RIB+AIB::HisCl (ii) based on time-varying (a) and time-invariant module assignments (b). While modularity decreases and becomes less stable for WT when calculated with time-invariant module assignments **(b, i)**, stronger decreases and destabilization are observed for AVB+RIB+AIB::HisCl. **(c, d)** Quantifying absolute (c) and relative (d) decrease in modularity (Q_TV_) after recalculating with time invariant module assignments (Q_TI_) reveals destabilization of module assignments after silencing several key hub interneurons. Neurons are ordered from left to right according to order listed in legend. Significance was tested using pairwise Welch’s t-tests between HisCl and WT with Bonferroni correction for multiple comparisons (*p<0.05, **p<0.01, ***p<0.001). **(e)** While silencing different key hub neurons has relatively little effect on stability of modularity (PSD max, x-axis), destabilization of module assignments (Q_TI_/Q_TV_, y-axis) is more variable across neurons.

To verify this, we quantified destabilization of module assignments by calculating both the decrease in modularity observed (both absolute and relative) when using time-invariant module assignments (Figure 6c, d). Some decrease in modularity is expected when switching from time-varying to time-invariant module assignments, as module assignments are no longer optimized for individual windows. However, more extensive reorganization across time will result in more extensive decreases in modularity, indicating increased instability of module assignments. Wild-type module assignments are relatively stable, resulting in relatively little decrease in modularity when calculated with time-invariant modules, compared to the decreases observed when silencing key hub neurons (Figure 6c, d). While AVB+RIB+AIB was the set of neurons that significantly destabilized modularity (Figure 5d), it is not the only set that significantly destabilizes module assignments. Silencing AVB and RIM individually and joint silencing of RIM+PVC and AVA+AVE+RIM+PVC also significantly destabilized module assignments (Figure 6c, d). Individually silencing AVE and RIB may also destabilize module assignments, but only relative decrease in modularity is significantly different from wild-type (Figure 6d). The effect of different neurons on stability of module assignments is much more varied than their effects on stability of modularity (Figure 6e). Comparison of time-varying and time-invariant module assignments for this dataset identified additional neurons that destabilize organization of time-varying functional connectivity when silenced. In the context of spontaneous, motor state encoding whole-brain activity, these neurons may serve to stabilize module membership over time or control stability, allowing reorganization in specific contexts.

## Discussion

In this work, we demonstrate the efficacy of modularity-based community detection for summarizing complex changes in time-varying functional connectivity. The compact summary provided by module assignments and modularity facilitates comparison of changes in functional connectivity across many time windows, individual recordings, and conditions. While modularity provides an initial readout of changes in time-varying functional connectivity, it does not fully reveal their nature: reorganization of network modules versus altered connectivity between stable modules. To assess this, we compare results obtained with time-varying and time-invariant module assignments. In principle, if similar results are observed with time-varying and time-invariant assignments, reorganization of modules is unlikely. Significant reorganization of modules should cause modularity (and observed trends) to change when calculated with time-invariant module assignments, instead of time-varying. We applied these analyses to both multi-cell food odor stimulus and spontaneous whole-brain activity datasets in this work. In multi-cell food odor data, we identified decreases in modularity during food odor stimulus within a small interneuron network (*glr-1p*) that did not rely on reorganization of network modules.

While modules were stable in response to food odor stimulus, we did find significant destabilization of modules in spontaneous whole-brain activity when several key hub interneurons were silenced (individual silencing of AVB and RIM and joint silencing of RIM+PVC, AVA+AVE+RIM+PVC, and AVB+RIB+AIB). Previously, these anatomical hubs were implicated in stabilizing pairwise correlations in an analysis of time-invariant functional connectivity.^17^ Our analysis of time-varying functional connectivity on the same dataset extends these findings by suggesting such anatomical hubs also stabilize modular organization of functional connectivity over time, in addition to stabilizing pairwise correlations. Interestingly, while some of the anatomical hubs tested significantly destabilized module assignments when silenced, not all of them had this effect. This suggests that while anatomical connectivity (and higher order topological features of the connectome) might be one mechanism nervous systems use to stabilize, control, and reorganize activity at intermediate scales (network modules), such organization cannot be explained by anatomy alone. These results call for further testing of what molecular mechanisms these anatomical hubs might use to stabilize modular organization of neural activity, control the extent of reorganization that is allowed, and on what timescales they might exert this control to enable multi-scale organization of neural activity.

While further investigation of the underlying molecular mechanisms that stabilize, control, and reorganize activity at intermediate scales is needed, it is of note that many neurons without direct anatomical synapses in *C. elegans* express complementary neuropeptide-receptor pairs.^50,51^ This prevalence of neuromodulatory volume transmission is not a unique feature of the *C. elegans* nervous system, but an important feature of many nervous systems, with relevance to many clinical disorders.^52^ This type of signaling might provide an additional mechanism through which nervous systems mediate intermediate-scale, modular organization of neural activity. Our results not only demonstrate the usefulness of modularity and module assignments for compactly summarizing changes in tine-varying functional connectivity, but also for connecting changes at brain-wide and single-cell scales to intermediate scale network modules. Making this connection is increasingly relevant in the context of technological advances that have increased the prevalence of large-scale, high resolution neural activity data across multiple model systems. When such recording methodologies are combined with experimental perturbation of potential molecular, anatomical, or functional mechanisms for organizing activity at intermediate scales, community detection and modularity provide a means for detecting changes at these scales.

## Methods

### *C. elegans* strains and maintenance

Multi-cell food odor stimulus and buffer control recordings were collected using *C. elegans* strain ZC3292 (*yxEx1701*[*glr-1p::GCaMP6s, glr-1p::NLS-mCherry-NLS*]) from previous work,^43^ which expresses nuclear-localized mCherry and cytosolic GCaMP6s in neurons that express the glutamate receptor subunit, *glr-1*(AVA, AVE, AVD, RMDD/V, RMD, RIM, SMDD/V, primarily locomotory command interneurons). This strain was maintained at 20 °C on NGM (3 g/L NaCl, 2.5 g/L peptone, 2% agar, 1 mM CaCl_2_, 5 mg/L cholesterol, 1 mM MgSO_4_, 25 mM KPO_4_ buffer) agar plates freshly seeded with *E. coli* OP50 (grown overnight at 37 °C in LB broth).^53^ For all imaging experiments, worms were well fed for at least 2 generations and age synchronized to Day 1 – 2 of adulthood using a timed-hatch off procedure, where adults and larvae were removed from a mixed population plate by rinsing with 1x S-basal buffer, and freshly hatched L1s were collected after allowing eggs to hatch 3 – 6 hours at 20 °C. The collected larvae were imaged 72 – 96 hours after hatching.

### Microfluidic device fabrication

Olfactory stimulation chips^38^ (72 um width x 24 um height) were fabricated using standard soft lithography methods.^54^ Briefly, PDMS elastomer was mixed in a 10:1 ratio of elastomer to cross-linker, poured onto an SU-8 master mold, and cured overnight at 75 °C. Cured PDMS was removed from the master mold, cut into individual devices, and inlets and outlets were punched using a 1mm biopsy punch. PDMS devices were bonded to #1.5 glass coverslips by treating both sides with air plasma for 30s and heating at 150 °C for at least 5 min.

### Multi-cell imaging with food odor stimulus

Food odor stimulus was prepared by inoculating 25 mL of LB broth in a 50 mL centrifuge tube with a single colony of E. coli OP50, shaking overnight at 37 °C (250 RPM), centrifuging (3,000 RCF for 10 min) to separate cell from supernatant, and adding 10 µM fluorescein dye to validate stimulus delivery. OP50 supernatant was used on the same day that it was prepared. All other device inlets were connected to reservoirs containing NGM Buffer (1mM CaCl_2_, 1 mM MgSO_4_, 25 mM KPO_4_ buffer). For the buffer-buffer control experiments, the fluorescein dyed OP50 supernatant stream was substituted with a fluorescein dyed NGM stream.

Volumetric, two-channel time series microscopy videos were collected using a Bruker Opterra II swept field microscope equipped with a 40x/0.75 air objective (Plan Fluor) and Photometrics Evolve 512 Delta EM-CCD. Excitation was alternated between 561 nm and 488 nm for each frame to image mCherry and GCaMP respectively and a 405 nm quad bandpass filter was used for emission. We obtained a volumetric imaging rate of ∼3.3 volumes/s (0.3024 s per volume) by adjusting Z-slice thickness (1.5 – 1.8 µm) to obtain a fixed number of 10 slices per volume and an exposure time of 15 ms per slice per channel.

During the recording, stimulus delivery was controlled using solenoid valves and a custom pressure control apparatus described previously^55^ with an updated MATLAB GUI. This updated GUI simultaneously triggers the start of stimulus delivery and the start of video acquisition using the Bruker Prairie View API. The stimulus sequence delivered consisted of 60 s of NGM buffer, before alternating between 30 s of OP50 and 60 s of NGM for four cycles (420 s total recording length, 1400 volumes). For buffer-buffer recordings, flow followed the same sequence, but switched between two NGM buffer streams, rather than buffer and OP50 supernatant.

### Multi-cell data pre-processing

The Bruker Opterra II provides recordings as a series of individual .ome.tif images. These were converted into a 5D (xyz, time, channel) .h5 using a custom MATLAB script (included on github). The individual neurons were tracked across time in the mCherry channel using ZephIR ^12^. Tracking results from ZephIR were manually inspected for accuracy and manually corrected tracks were used to extract fluorescence intensity of individual neurons over time with ZephIR’s built-in extract_traces function.

For each neuron, the ratio of GCaMP/mCherry fluorescence was taken to control for potential motion artifacts in the neural activity signal, *R*(*t*) = *F_GCaMP_*(*t*)/*F_mC_*_ℎ*erry*_(*t*). During tracking, some timepoints with high shear across the image volume (motion artifact) were identified and excluded from trace extraction. Signals for these missing timepoints were imputed using a 3 timepoint sliding window median filter (for single missing frames) and linear interpolation for larger gaps (max gap length = 3/1400 timepoints).

Recordings were split into individual trials of stimulus application (60 s pre, 30 s peri, 30s post stimulus activity, 120 s/397 volumes total) before baseline normalization of ratio by the signal 30 s before stimulus onset, *R_o_* = *mean*(*R*(30 *to* 60 *s*)), Δ*R*/*R_o_* = *R*(*t*) − *R_o_*.

### Spontaneous whole-brain data preprocessing

Spontaneous whole-brain neural activity data from Uzel et. al. 2022^17^ was downloaded from the OSF repository. Individual .mat files for different histamine silencing conditions were concatenated into a single array containing all histamine silencing conditions to facilitate comparison of different conditions.

### Community detection and calculation of network modularity

We calculated sliding window correlations between pairs of neurons to obtain time-varying functional connectivity. For the multi-cell food odor stimulus data, sliding window length was ∼20 s (66 timepoints) with 2 s offsets (6 timepoints). For whole-brain imaging data, sliding window length was 100 s with 10 s offsets. Rather than perform community detection (Louvain optimization of module assignments) directly on the sliding window correlations, we zeroed out the diagonal of these correlation matrices and then applied the Fisher transformation, *Z*(*t*) = arctanh(*A*(*t*)).^34,35^

For single-slice community detection, we applied Louvain optimization^56,57^ to each sliding window individually and used the following formulation of modularity, which accounts for correlation signs^46^ by splitting positive and negative Fisher transformed correlations 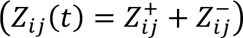. The equation below comparing observed correlation strength between a pair of neurons *ij* to average strength of positive/negative correlations for individual neurons 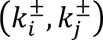 and overall strength of positive/negative correlations across all neurons (*m*^±^). In the equation below, *δ* is the Kronecker delta function and *g_i_*, *g*_j_are the module assignments of neurons *i* and *j*.

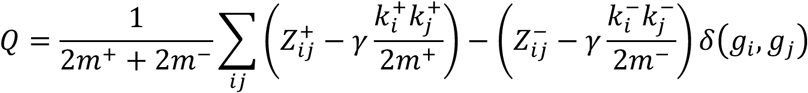

For multi-slice community detection, we used the formulation of modularity below^47^, which links adjacent timepoints with a coupling parameter, *ω*, which we varied from 0.01 to 100 and a fixed resolution parameter *γ* of 1. In the equation below, *i* and *j* are neuron indices, *s* and *r* are sliding window indices, *δ* is the Kronecker delta function, *g_is_* is the module assignment of neuron *i* in window *s*, and *μ* is a normalization constant of all possible contribution to modularity from 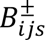 terms and coupling across windows, *ω*. The 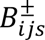 terms are the contributions of individual neuron pairs to modularity for window *s* and incorporate a modification of modularity that accounts for correlation signs.^46^

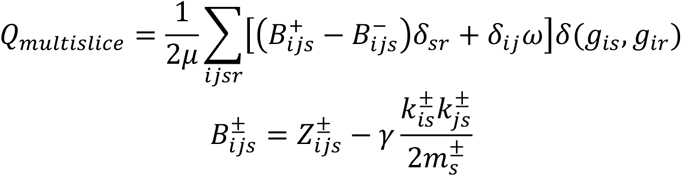

For both single-slice and multi-slice community detection, we repeated Louvain optimization and found a consensus solution across optimization runs^34,58^. For single-slice, we repeated optimization 100x (per sliding window). For multi-slice, we repeated optimization 1,000x and followed additional guidelines from Basset et. al^34^ by thresholding the consensus matrix by a shuffled null control. For individual windows, module assignments were re-indexed to minimize the number of neurons that swap modules between adjacent timepoints. After obtaining representative module assignments, we calculated modularity as a network metric, using *A*(*t*) directly (without zeroing out the diagonal or Fisher transforming) and the module assignments for each time point using equation below (same as previous, but with non-Fisher transformed correlations).

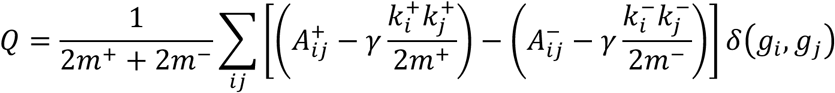

### Quantification of network-level responses to food odor stimulus

Network-level responses to food odor stimulus were quantified by subtracting the average modularity during buffer application from the average modularity during stimulus application. For this calculation, sliding windows were indexed to the last/end timepoint, aligning the start of stimulus application with the first window containing a stimulus-on.

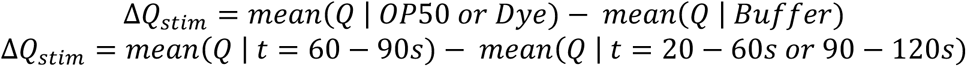

### Quantification of destabilization of network modularity in whole-brain activity

To quantify destabilization of network modularity in spontaneous whole-brain activity data, we calculate variance, max power spectral density, and autocorrelation of the modularity time series, *Q*. The variance is also equivalent to the total power spectral density (area under the curve of power spectra density versus frequency) of the modularity time series. The max power spectral density is the amount of variance that can be explained by a single sinusoidal frequency. Both will increase as modularity becomes less stable over time. From the autocorrelation, we obtain two metrics, the decorrelation time (DT), which is the first zero-crossing of the autocorrelation function, and the integrated decorrelation time (IDT), which is the area under the curve of the autocorrelation function. Both will increase as modularity becomes less stable over time. Equations for these metrics are provided below

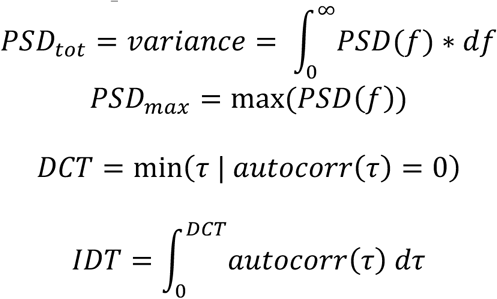

### Conversion of time-varying module assignments to time-invariant

Time varying module assignments were converted to time-invariant assignments by finding an allegiance matrix of module assignments across timepoints (probabilities of module co-assignment between neurons across time), then clustering the allegiance matrix based on weighted modularity and Louvain clustering. The time-invariant assignments were a consensus across 100 runs of Louvain optimization. This time-invariant module assignment was then used for calculation of modularity at each timepoint, instead of the time-varying module assignment.

### Joint phase randomization of functional connectivity

We phase randomized functional connectivity in figure 3 by taking the discrete Fourier transform of Fisher transformed sliding window correlations, preserving the Fourier amplitudes, randomizing the Fourier phases, and taking the inverse Fourier transform. We used the same set of random phases across all neuron pairs in a recording (joint phase randomization), rather than randomize pairs individually (individual phase randomization), to preserve covariance of sliding window correlations across neuron pairs.

### Community detection and calculation of modularity without correlation sign

To test the effect of community detection and calculation of modularity without accounting for correlation signs, we repeated the single-slice community detection procedure on the multi-cell food odor stimulus data, but took the absolute value of *A*(*t*)/*Z*(*t*) when calculating modularity and used the original formulation of modularity for weighted, undirected networks.^23^

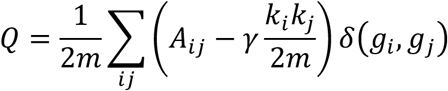

Aside from taking absolute value of correlations and using the modularity formula above, our approach was unchanged for this analysis. Community detection was performed using *Z*(*t*) with the diagonal zeroed out and a resolution parameter of 1. Representative module assignments for each timepoint were obtained by taking a consensus across 100 optimization runs.

Representative module assignments and untransformed correlation (*A*(*t*) including ones along the diagonal) were used to calculate modularity for individual timepoints.

### Statistical methods

All statistics and analysis were performed in MATLAB (version 2021b) with methods used for specific figures described below and test results provided as supplementary tables 1 – 6.

For figure 2h, significance of changes in Δ*Q_stim_* between OP50 and Buffer conditions were tested by fitting a linear mixed effect (LME) model with the form below. This model accounts for repeated observation of worms (4 trials per worm) and controls for potential trends across trials when comparing conditions. Stimulus was modeled as a categorical predictor and trial numbers were modeled as quantitative predictors. The data contained 5 worms per condition and 4 trials per worm (40 observations total).

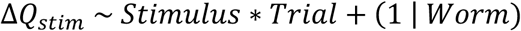

For figures 3d, 4b, and 4d, we tested the effect of different module assignment and modularity calculation methods on Δ*Q_stim_* by fitting expanded LME models, which compared values obtained using different methods by modeling method as an additional categorical predictor. For 3d, method corresponded to time-varying versus time-invariant modules. For 4b, method corresponded to correlation signs versus absolute value. For 4d, method corresponded to single-slice or multi-slice with different parameter values (multi-slice parameter values were modeled as categorial predictors, rather than quantitative).

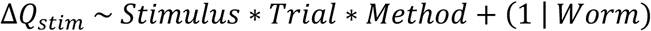

For these models in figures 3d, 4b, and 4d, the reference method was always single-slice module assignment with correlation signs considered (i.e. the results shown in figure 2h). After fitting the LME, pairwise contrasts between OP50 and Buffer stimulus were calculated for non-reference methods to obtain significance values. For 4d, p-values were adjusted using the Bonferroni correction to account for multiple-comparisons.

For figure 3e, the p-value reported is the fraction of phase randomized datasets (n = 1,000) where average Δ*Q_stim_* across all buffer trials is less than the average Δ*Q_stim_* across all OP50 trials.

For figure 4e, we subtracted Δ*Q_stim_* calculated using the single-slice method from Δ*Q_stim_* calculated with the multi-slice method (at different parameter values) for each trial. We tested the effect of multi-slice coupling parameter on difference in results between single and multi-cell by fitting the following LME model. We modeled coupling parameter as a categorical predictor, set the reference level as coupling parameter = 1, and calculated pairwise contrasts between observed deviation and zero for non-reference levels. After calculating pairwise contrasts, the p-values in 4e were adjusted using the Bonferroni correction to account for multiple-comparisons.

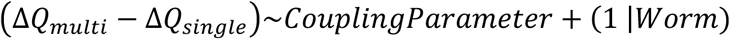

For figures 5d, 6c, and 6d, significance was tested using pairwise Welch’s t-tests between wild-type (WT) and different interneuron histamine silencing conditions. Bonferroni correction was applied to all p-values to account for multiple comparisons. This approach was taken to account for the imbalance in sample sizes between wild-type (n = 6) and histamine silencing recordings (n = 48 across 10 silencing conditions) in the overall dataset, but balanced sample sizes between wild-type and individual histamine silencing conditions (n = 4 – 6).

## Data and code availability

All data generated for this study will be made publicly available at an OSF repository upon acceptance. All analysis scripts will be made available at a GitHub repository upon acceptance.

## Acknowledgements

We thank Y. Moon, H. Matsumura, and G. Aubry for their comments on the manuscript. H.L and Y.Z are funded by NIH (MH130064 and NS115484).

## Competing Interests

The authors declare no competing interests.

## Notes

### Competing Interest Statement

The authors have declared no competing interest.

### Summary of Updates

Corrected acknowledgements, work was funded by MH130064, not MH117386.

## References

1 Steinmetz, N. A. et al. Neuropixels 2.0: A miniaturized high-density probe for stable, long-term brain recordings. Science 372 (2021). 10.1126/science.abf4588

2 Consortium, T. M. et al. Functional connectomics spanning multiple areas of mouse visual cortex. Nature 640, 435–447 (2025). 10.1038/s41586-025-08790-w

3 Ahrens, M. B., Orger, M. B., Robson, D. N., Li, J. M. & Keller, P. J. Whole-brain functional imaging at cellular resolution using light-sheet microscopy. Nature Methods 10, 413–420 (2013). 10.1038/nmeth.2434

4 Schrödel, T., Prevedel, R., Aumayr, K., Zimmer, M. & Vaziri, A. Brain-wide 3D imaging of neuronal activity in Caenorhabditis elegans with sculpted light. Nature Methods 10, 1013–1020 (2013). 10.1038/nmeth.2637

5 Lemon, W. C. et al. Whole-central nervous system functional imaging in larval Drosophila. Nature Communications 6, 7924 (2015). 10.1038/ncomms8924

6 Nguyen, J. P. et al. Whole-brain calcium imaging with cellular resolution in freely behaving Caenorhabditis elegans. Proceedings of the National Academy of Sciences 113, E1074–E1081 (2016). 10.1073/pnas.1507110112

7 Venkatachalam, V. et al. Pan-neuronal imaging in roaming Caenorhabditis elegans. Proceedings of the National Academy of Sciences 113, E1082–E1088 (2016). 10.1073/pnas.1507109113

8 Kim, D. H. et al. Pan-neuronal calcium imaging with cellular resolution in freely swimming zebrafish. Nature Methods 14, 1107–1114 (2017). 10.1038/nmeth.4429

9 Voleti, V. et al. Real-time volumetric microscopy of in vivo dynamics and large-scale samples with SCAPE 2.0. Nature Methods 16, 1054–1062 (2019). 10.1038/s41592-019-0579-4

10 Atanas, A. A. et al. Deep Neural Networks to Register and Annotate the Cells of the C. elegans Nervous System. bioRxiv, 2024.2007.2018.601886 (2024). 10.1101/2024.07.18.601886

11 Park, C. F. et al. Automated neuron tracking inside moving and deforming C. elegans using deep learning and targeted augmentation. Nature Methods 21, 142–149 (2024). 10.1038/s41592-023-02096-3

12 Ryu, J. et al. Versatile multiple object tracking in sparse 2D/3D videos via deformable image registration. PLOS Comput. Biol. 20, e1012075 (2024). 10.1371/journal.pcbi.1012075

13 Calhoun, W. A., Moon, S., Peng, L. & Lu, H. High speed functional imaging with a microfluidics-compatible open-top light-sheet microscope enabled by model predictive control of a tunable lens. bioRxiv, 2025.2007.2023.666439 (2025). 10.1101/2025.07.23.666439

14 Han, H., Pritchard, R. & Lu, H. ASCENT: Annotation-free Self-supervised Contrastive Embeddings for 3D Neuron Tracking in Fluorescence Microscopy. bioRxiv, 2025.2007.2023.666425 (2025). 10.1101/2025.07.23.666425

15 Ji, N. et al. A neural circuit for flexible control of persistent behavioral states. Elife 10 (2021). 10.7554/elife.62889

16 Yemini, E. et al. NeuroPAL: A Multicolor Atlas for Whole-Brain Neuronal Identification in C. elegans. Cell 184, 272–288.e211 (2021). 10.1016/j.cell.2020.12.012

17 Uzel, K., Kato, S. & Zimmer, M. A set of hub neurons and non-local connectivity features support global brain dynamics in C. elegans. Curr Biol (2022). 10.1016/j.cub.2022.06.039

18 Wirak, G. S., Florman, J., Alkema, M. J., Connor, C. W. & Gabel, C. V. Age-associated changes to neuronal dynamics involve a disruption of excitatory/inhibitory balance in C. elegans. Elife 11, e72135 (2022). 10.7554/elife.72135

19 Dag, U. et al. Dissecting the functional organization of the C. elegans serotonergic system at whole-brain scale. Cell 186, 2574–2592.e2520 (2023). 10.1016/j.cell.2023.04.023

20 Randi, F., Sharma, A. K., Dvali, S. & Leifer, A. M. Neural signal propagation atlas of Caenorhabditis elegans. Nature 623, 406–414 (2023). 10.1038/s41586-023-06683-4

21 Kramer, T. S. et al. Neural Sequences Underlying Directed Turning in C. elegans. bioRxiv, 2024.2008.2011.607076 (2024). 10.1101/2024.08.11.607076

22 Fieseler, C. et al. An intrinsic neuronal manifold underlies brain-wide hierarchical organization of behavior in C. elegans. bioRxiv, 2025.2003.2009.642241 (2025). 10.1101/2025.03.09.642241

23 Newman, M. E. J. Modularity and community structure in networks. Proceedings of the National Academy of Sciences 103, 8577–8582 (2006). 10.1073/pnas.0601602103

24 Fortunato, S. Community detection in graphs. Phys. Rep. 486, 75–174 (2010). 10.1016/j.physrep.2009.11.002

25 Fortunato, S. & Hric, D. Community detection in networks: A user guide. Phys. Rep. 659, 1–44 (2016). 10.1016/j.physrep.2016.09.002

26 Tóth, G. et al. Inequality is rising where social network segregation interacts with urban topology. Nature Communications 12, 1143 (2021). 10.1038/s41467-021-21465-0

27 Wu, X. et al. Selective enrichment of bacterial pathogens by microplastic biofilm. Water Res. 165, 114979 (2019). 10.1016/j.watres.2019.114979

28 Greenblum, S., Turnbaugh, P. J. & Borenstein, E. Metagenomic systems biology of the human gut microbiome reveals topological shifts associated with obesity and inflammatory bowel disease. Proceedings of the National Academy of Sciences 109, 594–599 (2012). 10.1073/pnas.1116053109

29 Wiedmer, R. & Griffis, S. E. Structural characteristics of complex supply chain networks. J. Bus. Logist. 42, 264–290 (2021). 10.1111/jbl.12283

30 Xu, M., Pan, Q., Muscoloni, A., Xia, H. & Cannistraci, C. V. Modular gateway-ness connectivity and structural core organization in maritime network science. Nature Communications 11, 2849 (2020). 10.1038/s41467-020-16619-5

31 Hagmann, P. et al. Mapping the Structural Core of Human Cerebral Cortex. PLoS Biol. 6, e159 (2008). 10.1371/journal.pbio.0060159

32 Fair, D. A. et al. Functional Brain Networks Develop from a “Local to Distributed” Organization. PLOS Comput. Biol. 5, e1000381 (2009). 10.1371/journal.pcbi.1000381

33 Bassett, D. S. et al. Dynamic reconfiguration of human brain networks during learning. Proceedings of the National Academy of Sciences 108, 7641–7646 (2011). 10.1073/pnas.1018985108

34 Bassett, D. S. et al. Robust detection of dynamic community structure in networks. Chaos: Interdiscip. J. Nonlinear Sci. 23, 013142 (2013). 10.1063/1.4790830

35 Bassett, D. S., Yang, M., Wymbs, N. F. & Grafton, S. T. Learning-induced autonomy of sensorimotor systems. Nat. Neurosci. 18, 744–751 (2015). 10.1038/nn.3993

36 Sporns, O. & Betzel, R. F. Modular Brain Networks. Annu. Rev. Psychol. 67, 1–28 (2015). 10.1146/annurev-psych-122414-033634

37 Gordus, A., Pokala, N., Levy, S., Flavell, S. W. & Bargmann, C. I. Feedback from network states generates variability in a probabilistic olfactory circuit. Cell 161, 215 227 (2015). 10.1016/j.cell.2015.02.018

38 Chronis, N., Zimmer, M. & Bargmann, C. I. Microfluidics for in vivo imaging of neuronal and behavioral activity in Caenorhabditis elegans. Nature Methods 4, 727–731 (2007). 10.1038/nmeth1075

39 Lee, H. J., Vallier, J. & Lu, H. Microfluidic localized hydrogel polymerization enables simultaneous recording of neural activity and behavior in C. elegans. React. Chem. Eng. 9, 666–676 (2023). 10.1039/d3re00516j

40 Seyedolmohadesin, M. et al. Whole-brain chemosensory responses of both C. elegans sexes. bioRxiv, 2025.2005.2015.654129 (2025). 10.1101/2025.05.15.654129

41 Susoy, V. et al. Natural sensory context drives diverse brain-wide activity during C. elegans mating. Cell 184, 5122–5137.e5117 (2021). 10.1016/j.cell.2021.08.024

42 Atanas, A. A. et al. Brain-wide representations of behavior spanning multiple timescales and states in C. elegans. Cell 186, 4134–4151.e4131 (2023). 10.1016/j.cell.2023.07.035

43 Lee, H. J. et al. Automated cell annotation in multi-cell images using an improved CRF_ID algorithm. Elife 12, RP89050 (2025). 10.7554/elife.89050

44 Kato, S. et al. Global Brain Dynamics Embed the Motor Command Sequence of Caenorhabditis elegans. Cell 163, 656–669 (2015). 10.1016/j.cell.2015.09.034

45 Good, B. H., Montjoye, Y.-A. d. & Clauset, A. Performance of modularity maximization in practical contexts. Phys. Rev. E 81, 046106 (2010). 10.1103/physreve.81.046106

46 Gómez, S., Jensen, P. & Arenas, A. Analysis of community structure in networks of correlated data. Phys. Rev. E 80, 016114 (2009). 10.1103/physreve.80.016114

47 Mucha, P. J., Richardson, T., Macon, K., Porter, M. A. & Onnela, J.-P. Community Structure in Time-Dependent, Multiscale, and Multiplex Networks. Science 328, 876–878 (2010). 10.1126/science.1184819

48 Honey, C. J. et al. Predicting human resting-state functional connectivity from structural connectivity. Proceedings of the National Academy of Sciences 106, 2035–2040 (2009). 10.1073/pnas.0811168106

49 Pokala, N., Liu, Q., Gordus, A. & Bargmann, C. I. Inducible and titratable silencing of Caenorhabditis elegans neurons in vivo with histamine-gated chloride channels. Proceedings of the National Academy of Sciences of the United States of America 111, 2770–2775 (2014). 10.1073/pnas.1400615111

50 Beets, I. et al. System-wide mapping of peptide-GPCR interactions in C. elegans. Cell Rep. 42, 113058 (2023). 10.1016/j.celrep.2023.113058

51 Ripoll-Sánchez, L. et al. The neuropeptidergic connectome of C. elegans. Neuron 111, 3570–3589.e3575 (2023). 10.1016/j.neuron.2023.09.043

52 Özçete, Ö. D., Banerjee, A. & Kaeser, P. S. Mechanisms of neuromodulatory volume transmission. Mol. Psychiatry 29, 3680–3693 (2024). 10.1038/s41380-024-02608-3

53 Stiernagle, T. in WormBook (ed The C. elegans Research Community) (WormBook, 2006).

54 San-Miguel, A. & Lu, H. in WormBook (ed The C. elegans Research Community) (WormBook, 2013).

55 San-Miguel, A. et al. Deep phenotyping unveils hidden traits and genetic relations in subtle mutants. Nature Communications 7, 12990 (2016). 10.1038/ncomms12990

56 Blondel, V. D., Guillaume, J.-L., Lambiotte, R. & Lefebvre, E. Fast unfolding of communities in large networks. J. Stat. Mech.: Theory Exp. 2008, P10008 (2008). 10.1088/1742-5468/2008/10/p10008

57 Jeub, L. G. S., Bazzi, M., Jutla, I. S. & Mucha, P. J. GenLouvain: A generalized Louvain method for community detection implemented in MATLAB. (2011).

58 Lancichinetti, A. & Fortunato, S. Consensus clustering in complex networks. Sci. Rep. 2, 336 (2012). 10.1038/srep00336

