## Supplementary material for "Network modularity reveals context and state-dependent reorganization of time-varying functional connectivity in single-cell resolved neural activity recordings": SI figures and tables

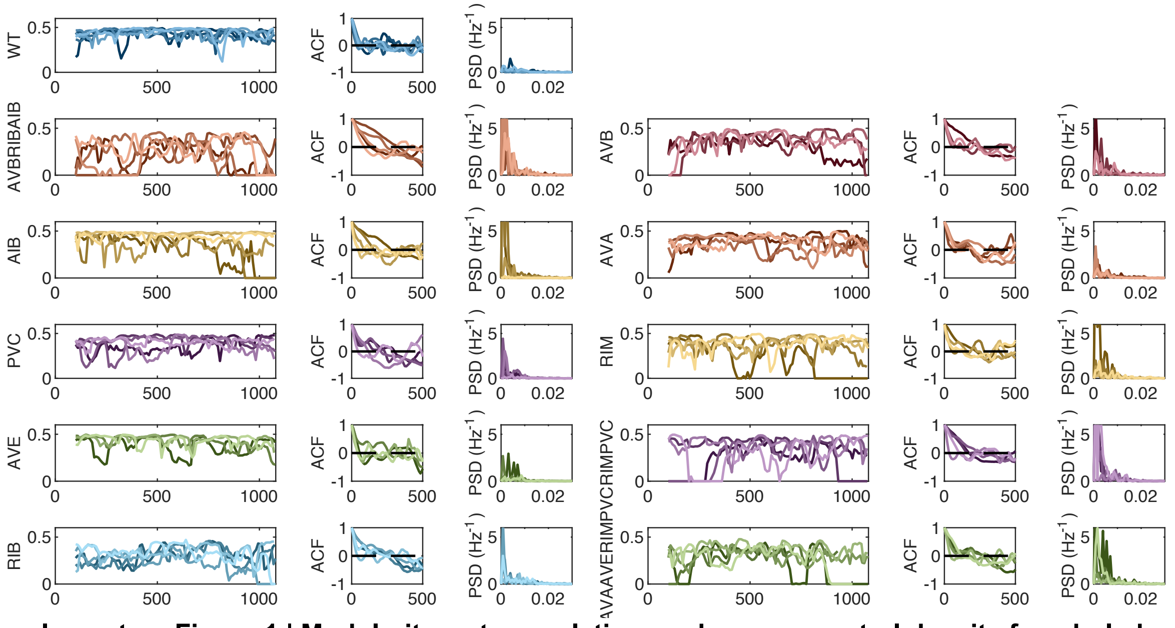


**Supplementary Figure 1 | Modularity, autocorrelation, and power spectral density for whole-brain data with multi-slice module assignment (coupling parameter = 0.01)**

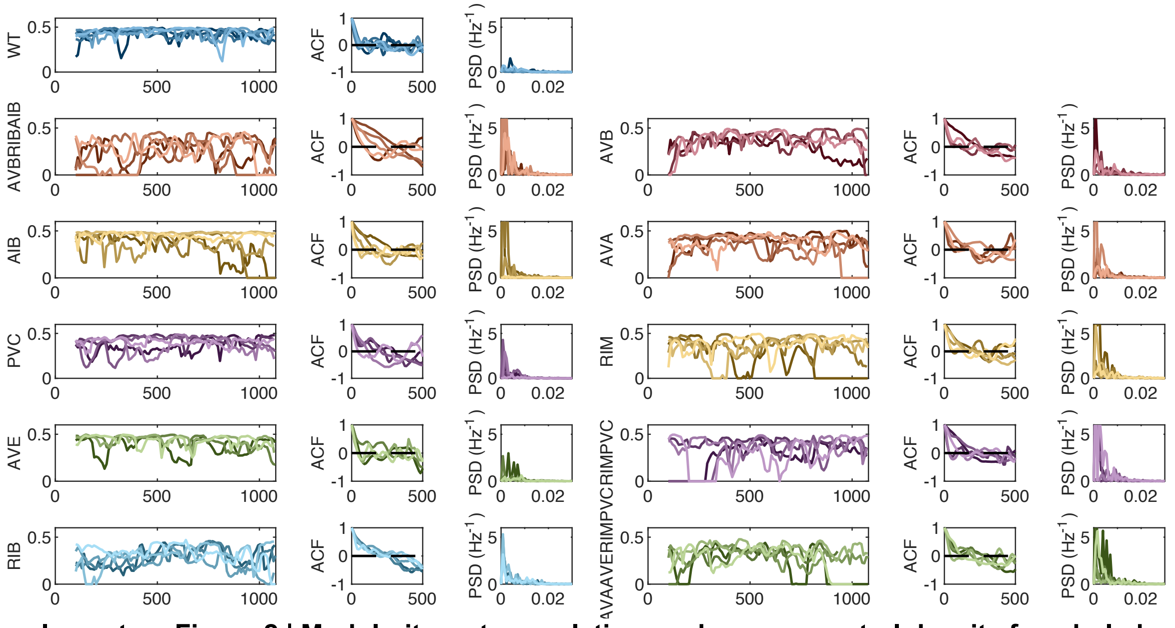


**Supplementary Figure 2 | Modularity, autocorrelation, and power spectral density for whole-brain data with multi-slice module assignment (coupling parameter = 0.1)**

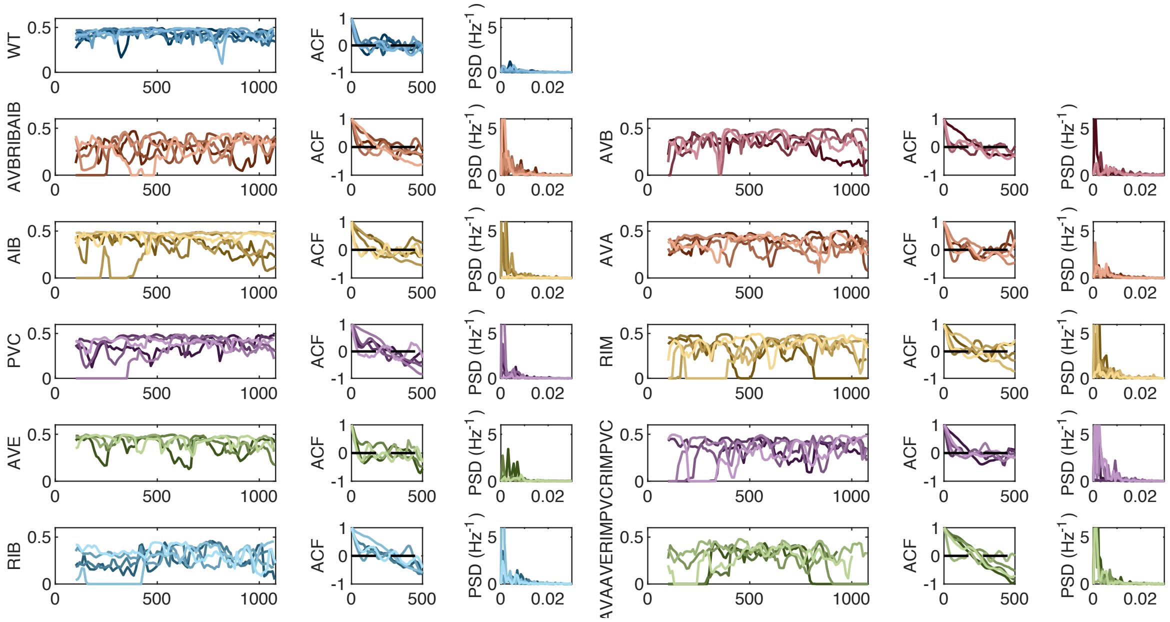


**Supplementary Figure 3 | Modularity, autocorrelation, and power spectral density for whole-brain data with multi-slice module assignment (coupling parameter = 1)**

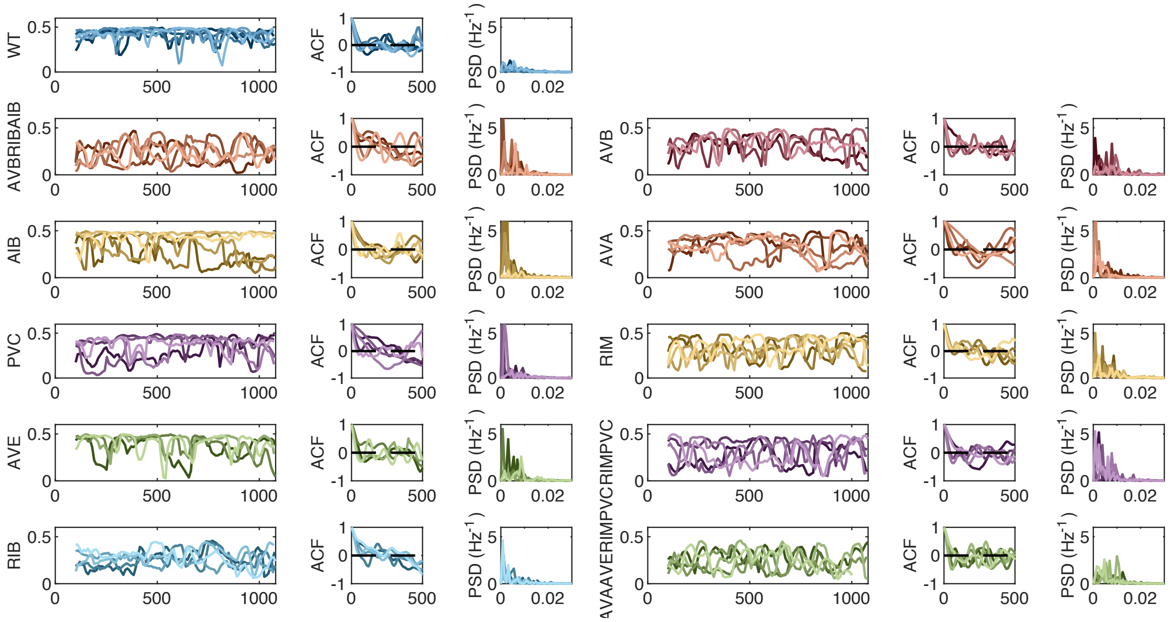


**Supplementary Figure 4 | Modularity, autocorrelation, and power spectral density for whole-brain data with multi-slice module assignment (coupling parameter = 10)**

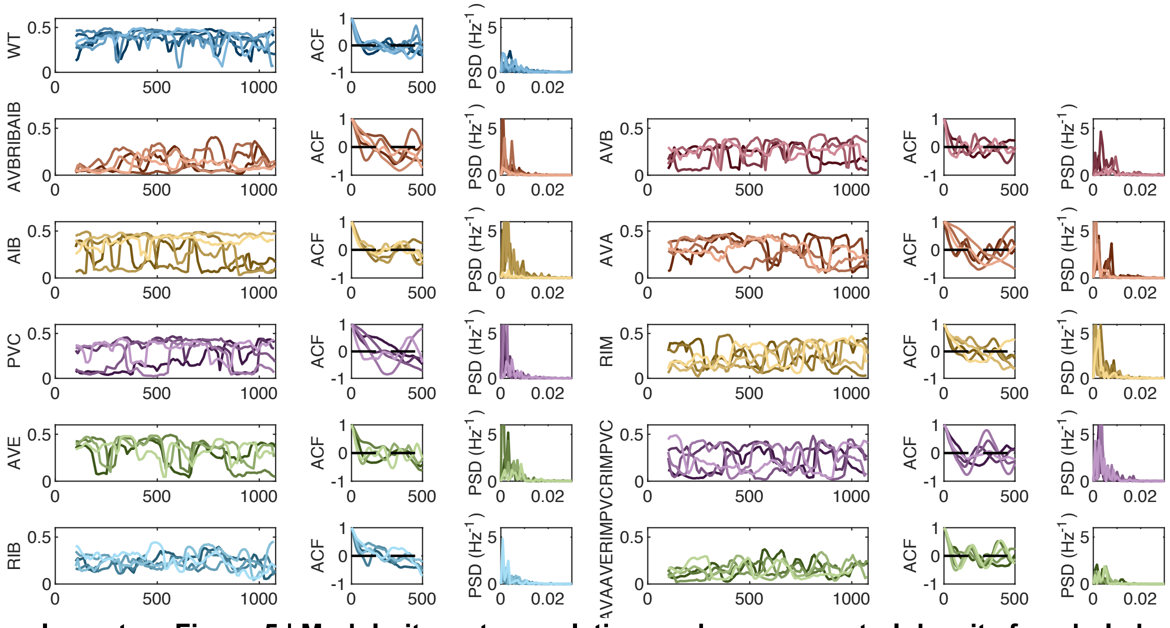


**Supplementary Figure 5 | Modularity, autocorrelation, and power spectral density for whole-brain data with multi-slice module assignment (coupling parameter = 100)**

**Supplementary Table 1 | Figure 2h LME model fit results**

| Model Information | Value |
|---|---|
| Fit method | REML |
| Number of observations | 40 |
| Fixed effects coefficients | 4 |
| Random effects coefficients | 10 |
| Covariance parameters | 2 |

**Model Fit Statistics**

| AIC | BIC | LogLikelihood | Deviance |
|---|---|---|---|
| -87.956 | -78.455 | 49.978 | -99.956 |

| Fixed Effects | Estimate | SE | DOF | t | p | 95% CI |
|---|---|---|---|---|---|---|
| (Intercept) | -0.077 | 0.019 | 36 | -4.068 | 0.00025 | [-0.116, -0.039] |
| Stimulus (Buffer) | 0.095 | 0.027 | 36 | 3.54 | 0.0011 | [0.041, 0.150] |
| Trial | 0.007 | 0.010 | 36 | 0.735 | 0.47 | [-0.013, 0.028] |
| Stimulus × Trial | -0.015 | 0.014 | 36 | -1.066 | 0.29 | [-0.045, 0.014] |

| Random Effects | Estimate (SD) | 95% CI |
|---|---|---|
| Worm (10 levels, Intercept SD) | 0 | [NaN, NaN] |
| Residual (Error SD) | 0.051 | [0.040, 0.064] |

**Supplementary Table 2 | Figure 3d LME model fit results**

| Model Information | Value |
| --- | --- |
| Fit method | REML |
| Number of observations | 80 |
| Fixed effects coefficients | 8 |
| Random effects coefficients | 10 |
| Covariance parameters | 2 |

**Model Fit Statistics**

| AIC | BIC | LogLikelihood | Deviance |
| --- | --- | --- | --- |
| -165.52 | -142.75 | 92.758 | -185.52 |

| Fixed Effects | Estimate | SE | DOF | t | p | 95% CI |
| --- | --- | --- | --- | --- | --- | --- |
| (Intercept) | -0.077 | 0.021 | 72 | -3.681 | 0.00045 | [-0.119, -0.035] |
| Stimulus (Buffer) | 0.095 | 0.030 | 72 | 3.203 | 0.0020 | [0.036, 0.154] |
| Trial | 0.007 | 0.011 | 72 | 0.665 | 0.51 | [-0.015, 0.030] |
| Method (invariant) | 0.016 | 0.030 | 72 | 0.522 | 0.60 | [-0.044, 0.075] |
| Stimulus × Trial | -0.015 | 0.016 | 72 | -0.965 | 0.34 | [-0.047, 0.016] |
| Stimulus × Method | -0.008 | 0.042 | 72 | -0.181 | 0.86 | [-0.091, 0.076] |
| Trial × Method | 0.001 | 0.016 | 72 | 0.0602 | 0.95 | [-0.031, 0.033] |
| Stimulus × Trial × Method | 0.002 | 0.023 | 72 | 0.107 | 0.92 | [-0.042, 0.047] |

| Random Effects | Estimate (SD) | 95% CI |
| --- | --- | --- |
| Worm (10 levels, Intercept SD) | 0 | [NaN, NaN] |
| Residual (Error SD) | 0.056 | [0.048, 0.066] |

| Additional Contrasts | p |
| --- | --- |
| OP50 vs. Buffer for Method (invariant) | 0.0043 |

**Supplementary Table 3 | Figure 4b LME model fit results**

| Model Information | Value |
|---|---|
| Fit method | REML |
| Number of observations | 80 |
| Fixed effects coefficients | 8 |
| Random effects coefficients | 10 |
| Covariance parameters | 2 |

**Model Fit Statistics**

| AIC | BIC | LogLikelihood | Deviance |
|---|---|---|---|
| -215.64 | -192.87 | 117.82 | -235.64 |

| Fixed Effects | Estimate | SE | DOF | t | p | 95% CI |
|---|---|---|---|---|---|---|
| (Intercept) | -0.077 | 0.015 | 72 | -5.213 | 1.7E-06 | [-0.107, -0.048] |
| Stimulus (Buffer) | 0.095 | 0.021 | 72 | 4.536 | 2.2E-05 | [0.053, 0.137] |
| Trial | 0.007 | 0.008 | 72 | 0.942 | 0.35 | [-0.008, 0.023] |
| Method (Abs) | 0.056 | 0.021 | 72 | 2.685 | 0.0090 | [0.015, 0.098] |
| Stimulus × Trial | -0.015 | 0.011 | 72 | -1.366 | 0.18 | [-0.038, 0.007] |
| Stimulus × Method (Abs) | -0.067 | 0.030 | 72 | -2.257 | 0.027 | [-0.126, -0.008] |
| Trial × Method (Abs) | -0.004 | 0.011 | 72 | -0.367 | 0.71 | [-0.027, 0.018] |
| Stimulus × Trial × Method (Abs) | 0.009 | 0.016 | 72 | 0.555 | 0.58 | [-0.023, 0.040] |

| Random Effects | Estimate (SD) | 95% CI |
|---|---|---|
| Worm (10 levels, Intercept SD) | 0 | [NaN, NaN] |
| Residual (Error SD) | 0.040 | [0.034, 0.047] |

| Additional Contrasts | p |
|---|---|
| OP50 vs. Buffer for Method (Abs) | 0.18 |

**Supplementary Table 4 | Figure 4d LME model fit results**

| Model Information | Value |
|---|---|
| Fit method | REML |
| Number of observations | 240 |
| Fixed effects coefficients | 24 |
| Random effects coefficients | 10 |
| Covariance parameters | 2 |

**Model Fit Statistics**

| AIC | BIC | LogLikelihood | Deviance |
|---|---|---|---|
| -523.3 | -435.54 | 287.65 | -575.3 |

| Fixed Effects | Estimate | SE | DOF | t | p | 95% CI |
|---|---|---|---|---|---|---|
| (Intercept) | -0.077 | 0.022 | 216 | -3.565 | 0.00045 | [-0.120, -0.035] |
| Condition (Buffer) | 0.095 | 0.031 | 216 | 3.102 | 0.0022 | [0.035, 0.156] |
| Trial | 0.007 | 0.010 | 216 | 0.716 | 0.48 | [-0.013, 0.028] |
| Method_0.01 | 0.045 | 0.028 | 216 | 1.641 | 0.10 | [-0.009, 0.100] |
| Method_0.1 | 0.024 | 0.028 | 216 | 0.875 | 0.38 | [-0.030, 0.079] |
| Method_1 | 0.014 | 0.028 | 216 | 0.490 | 0.63 | [-0.041, 0.068] |
| Method_10 | 0.004 | 0.028 | 216 | 0.137 | 0.89 | [-0.051, 0.058] |
| Method_100 | 0.023 | 0.028 | 216 | 0.830 | 0.41 | [-0.032, 0.077] |
| Condition × Trial | -0.015 | 0.015 | 216 | -1.038 | 0.30 | [-0.044, 0.014] |
| Condition × Method_0.01 | -0.015 | 0.039 | 216 | -0.379 | 0.71 | [-0.092, 0.062] |
| Condition × Method_0.1 | -0.003 | 0.039 | 216 | -0.082 | 0.94 | [-0.080, 0.074] |
| Condition × Method_1 | 0.018 | 0.039 | 216 | 0.458 | 0.65 | [-0.059, 0.095] |
| Condition × Method_10 | 0.007 | 0.039 | 216 | 0.180 | 0.86 | [-0.070, 0.084] |
| Condition × Method_100 | -0.017 | 0.039 | 216 | -0.422 | 0.67 | [-0.094, 0.061] |
| Trial × Method_0.01 | -0.012 | 0.015 | 216 | -0.835 | 0.40 | [-0.041, 0.017] |
| Trial × Method_0.1 | -0.001 | 0.015 | 216 | -0.085 | 0.93 | [-0.030, 0.028] |
| Trial × Method_1 | 0.003 | 0.015 | 216 | 0.183 | 0.86 | [-0.026, 0.032] |
| Trial × Method_10 | 0.000 | 0.015 | 216 | 0.016 | 0.99 | [-0.029, 0.029] |
| Trial × Method_100 | 0.001 | 0.015 | 216 | 0.099 | 0.92 | [-0.028, 0.031] |
| Condition × Trial × Method_0.01 | 0.007 | 0.021 | 216 | 0.343 | 0.73 | [-0.034, 0.048] |
| Condition × Trial × Method_0.1 | 0.002 | 0.021 | 216 | 0.075 | 0.94 | [-0.040, 0.043] |
| Condition × Trial × Method_1 | -0.016 | 0.021 | 216 | -0.776 | 0.44 | [-0.057, 0.025] |
| Condition × Trial × Method_10 | -0.004 | 0.021 | 216 | -0.211 | 0.83 | [-0.046, 0.037] |
| Condition × Trial × Method_100 | 0.001 | 0.021 | 216 | 0.036 | 0.97 | [-0.040, 0.042] |

| Random Effects | Estimate (SD) | 95% CI |
|---|---|---|
| Worm (10 levels, Intercept SD) | 0.021 | [0.011, 0.039] |
| Residual (Error SD) | 0.052 | [0.047, 0.058] |

| Additional Contrasts | p | p_corrected |
|---|---|---|
| OP50 vs. Buffer for Method_0.01 | 0.0095 | 0.057 |
| OP50 vs. Buffer for Method_0.1 | 0.003 | 0.018 |
| OP50 vs. Buffer for Method_1 | 0.0003 | 0.0018 |
| OP50 vs. Buffer for Method_10 | 0.001 | 0.006 |
| OP50 vs. Buffer for Method_100 | 0.011 | 0.066 |

**Supplementary Table 5 | Figure 4e LME model fit results**

| Model Information | Column1 |
|---|---|
| Fit method | REML |
| Number of observations | 200 |
| Fixed effects coefficients | 5 |
| Random effects coefficients | 10 |
| Covariance parameters | 2 |

**Model Fit Statistics**

| AIC | BIC | LogLikelihood | Deviance |
|---|---|---|---|
| -678.81 | -655.9 | 346.4 | -692.81 |

| Fixed Effects | Estimate | SE | DOF | t | p | 95% CI |
|---|---|---|---|---|---|---|
| (Intercept) | 0.014 | 0.006 | 195 | 2.326 | 0.021 | [0.002, 0.027] |
| Parameter 0.1 | 0.007 | 0.009 | 195 | 0.856 | 0.39 | [-0.009, 0.025] |
| Parameter 0.01 | 0.010 | 0.009 | 195 | 1.194 | 0.23 | [-0.007, 0.028] |
| Parameter 10 | -0.010 | 0.009 | 195 | -1.148 | 0.25 | [-0.027, 0.007] |
| Parameter 100 | 0.003 | 0.009 | 195 | 0.351 | 0.73 | [-0.014, 0.020] |

| Ranodm Effects | Estimate (SD) | 95% CI |
|---|---|---|
| Worm (10 levels, Intercept SD) | 0 | [NaN, NaN] |
| Residual (Error SD) | 0.039 | [0.035, 0.043] |

| Additional Contrasts | p | p_corrected |
|---|---|---|
| OP50 vs. Buffer for Parameter_0.1 | 0.0005 | 0.0025 |
| OP50 vs. Buffer for Parameter_0.01 | 0.0001 | 0.0005 |
| OP50 vs. Buffer for Parameter_10 | 0.48 | 1.00 |
| OP50 vs. Buffer for Parameter_100 | 0.0053 | 0.027 |

**Supplementary Table 6 | Figures 5d, 6c, and 6d statistical testing results**

P values were obtained by pairwise Welch's t-tests between histamine silencing and wild-type data with Bonferroni correction for multiple comparisons (n = 10 tests)

| Silenced Neuron | # Recordings | PSD max | Variance | Integrated Decorrelation Time | Decorrelation Time | Q_TI - Q_TV | Q_TI /Q_TV |
|---|---|---|---|---|---|---|---|
| AVB+RIB+AIB | 5 | **0.029*** | **0.040*** | 0.17 | 0.51 | **0.0013*** | **0.0012*** |
| AIB | 5 | 1.00 | 1.00 | 1.00 | 1.00 | 1.00 | 1.00 |
| PVC | 5 | 0.34 | 1.00 | 0.30 | 0.44 | 0.38 | 0.34 |
| AVE | 4 | 1.00 | 1.00 | 1.00 | 1.00 | 0.056 | **0.024*** |
| RIB | 5 | 0.23 | 0.17 | 1.00 | 1.00 | 0.21 | **0.011*** |
| AVB | 4 | 0.93 | 0.80 | 1.00 | 1.00 | **0.021*** | **0.017*** |
| AVA | 5 | 0.29 | 0.47 | 1.00 | 1.00 | 1.00 | 0.90 |
| RIM | 5 | 0.67 | 0.84 | 0.51 | 0.42 | **0.018*** | **0.0060*** |
| RIM+PVC | 5 | 1.00 | 0.84 | 1.00 | 1.00 | **0.0092*** | **0.020*** |
| AVA+AVE+RIM+PVC | 5 | 0.89 | 1.00 | 1.00 | 1.00 | **0.0026*** | **<0.0001*** |
